# A collection of *Trichoderma* isolates from natural environments in Sardinia, a biodiversity hotspot, reveals a complex virome that includes negative-stranded mycoviruses with unprecedented genome organizations

**DOI:** 10.1101/2023.03.31.535183

**Authors:** Saul Pagnoni, Safa Oufensou, Virgilio Balmas, Daniela Bulgari, Emanuela Gobbi, Marco Forgia, Quirico Migheli, Massimo Turina

**Affiliations:** Department of Agricultural and Environmental Sciences – Production, Landscape, Agroenergy, University of Milan, via Celoria 2, 20133, Milan, Italy; Department of Agricultural Sciences and NRD – Desertification Research Center, University of Sassari, Viale Italia 39a, I-07100 Sassari, Italy; Department of Molecular and Translational Medicine, University of Brescia, via Branze 39, 25123, Brescia, Italy; Institute for Sustainable Plant Protection, National Research Council of Italy, Strada delle Cacce, 73, 10135, Torino, Italy

## Abstract

The *Trichoderma* genus includes soil-inhabiting fungi that provide important ecological services in their interaction with plants and other fungi. They are exploited for biocontrol. A collection of *Trichoderma* isolates from the Sardinia island (a biodiversity hotspot) had been previously characterized. Here we started a characterization of the viral components associated to 113 selected *Trichoderma* isolates, representatives of the collection. We carried out NGS sequencing of ribosome depleted total RNA following a bioinformatic pipeline that detects virus RNA-dependent RNA polymerases (RdRP) and other conserved virus protein sequences. This pipeline detected 17 viral RdRPs. Two of them correspond to viruses already detected in other regions of the world. The remaining 15 represent isolates of new virus species: surprisingly, eight of them are from new negative stranded RNA viruses, which for the first time are reported in the genus *Trichoderma*. Among them is a cogu-like virus, very closely related to plant-infecting viruses. Regarding the positive strand viruses, it is noticeable the presence of an ormycovirus belonging to a recently characterized group of bi-segmented ssRNA genome viruses with still uncertain phylogenetic assignment. Finally, for the first time we report a bipartite mononegavirales-infecting fungi: the proteins encoded by the second genomic RNA were used to re-evaluate a number of viruses in the *Penicillimonavirus* and *Plasmopamonavirus* genera, here shown to be bipartite and to encode a conserved polypeptide having structural conservation with the nucleocapsid (NC) domain of members of the Rabhdoviridae.

IMPORTANCE

*Trichoderma* is a genus of fungi of great biotechnological impact in multiple industrial fields. The possibility to investigate a diverse collection of *Trichoderma* isolates allowed us to characterize both double-stranded and single-stranded virus genomes belonging to three of the major phyla that constitute the RNA viral kingdom, thus further increasing the taxa of viruses infecting this genus. To our knowledge here we report for the first time negative-stranded RNA viruses infecting *Trichoderma* spp. and through *in silico* structural analysis a new conserved domain of nucleocapsids common among some mymonavirids. Obtaining such a library of mycoviruses could be the basis for further development of targeted virus-induced gene silencing or gene editing (VIGS/VIGE) tools; in addition, the many biotechnological applications of this fungus, will require to assess the qualitative (commercial) stability of strains, linked to positive or negative effects caused by mycovirus infections.

## INTRODUCTION

*Trichoderma* is a genus of fungi that includes many species commonly isolated from soil and rhizosphere, where they live as saprophytes playing a significant role in the degradation of plant polysaccharides. Beside their widely recognized ecological role these filamentous fungi are also the most used bio-fungicides and plant growth modifiers in agriculture, and are sources of enzymes of industrial utility, including those used in the biofuels industry. To understand the impact that *Trichoderma* strains have on agriculture, it suffices to say that more than 60% of world registered bio-fungicides are based on species from this genus (1). Furthermore, they are prolific producers of secondary metabolites, some having clinical significance (2, 3), while other species have been engineered to work as microbial cell factories for the heterologous production of important proteins (4). *Trichoderma* species are also used for bioremediation applications, due to their ability to degrade and/or mobilize both organic and inorganic waste compounds, including heavy metals (5) and for the biorefinery industry, contributing to valorise agricultural wastes and by-products (6). Currently, these fungi are among the most widely studied organisms, as evidenced by the wealth of published literature and the number of patents being registered (7).

Mycoviruses, defined as viruses infecting and replicating in true fungi (kingdom *Eumycota*) or oomycetes (kingdom *Chromista*, phylum *Heterokonta*), have been identified for the first time more than 50 years ago, but only during the last decade a considerable amount of knowledge has started to accumulate, thus contributing to shed light on this previously unexplored virosphere domain (8). Currently, according to the official “Master Species list” (version 2021_v3, published in November 2022) provided by the International Committee on Taxonomy of Viruses, mycoviruses are taxonomically classified within 25 different officially recognized families. The majority of them (included in 12 families) possess a positive sense RNA (+RNA) genome and are accommodated within the *Alphaflexiviridae*, *Barnaviridae*, *Botourmiaviridae*, *Deltaviridae*, *Endornaviridae*, *Fusariviridae*, *Gammaflexiviridae*, *Hadakaviridae*, *Hypoviridae*, *Mitoviridae*, *Narnaviridae* and *Yadokaviridae* families. Another considerable number of fungal viruses possess a double-stranded RNA (dsRNA) genome and belong to 8 families, namely: *Chrysoviridae*, *Curvulaviridae*, *Megabirnaviridae*, *Partitiviridae*, *Polymycoviridae*, *Quadriviridae*, *Spinareoviridae* and *Totiviridae*; while just 4 families accommodate mycoviruses possessing a negative sense RNA (-RNA) genome, which are: *Discoviridae*, *Mymonaviridae*, *Phenuiviridae* and *Tulasviridae*. Finally, just one family named *Genomoviridae*, includes viruses having a circular single-stranded DNA (ssDNA) genome. Despite the extensive research that, in latest years, led to the discovery of diverse fungal viruses, the majority of information available concerns phytopathogenic fungi, while relatively little is known on viruses infecting free-living soil fungi, endophytic fungi, and/or epiphytic fungi (8).

Given the ecological relevance and the biotechnological impact of the *Trichoderma* genus, it is somewhat surprising that their associated mycovirome was only recently investigated: the first studies in this context mainly identified mycoviruses possessing a dsRNA genome, some of which have been characterized at a molecular level by providing their complete genome sequence (9–15); while three of them were classified as members of the *Partitiviridae* family (10, 12, 15) and one as member of the *Curvulaviridae* family (13), the majority could not be assigned to any officially recognized viral family (unclassified dsRNA). More recently, also +RNA mycoviruses have been identified and characterized in *Trichoderma*, two belonging to the family *Hypovirida*e (16, 17) and one to the newly proposed family Ambiguiviridae (18). Curiously, all of these mycoviruses were observed in association with *Trichoderma* strains collected in the Asian continent (mainly in China or Korea), while, to our knowledge, the mycovirome associated with other continents’ populations of the fungus has never been explored.

In addition, only a few of the above-mentioned viruses were cured from the original strains or transferred to new virus-free isolates by anastomosis, therefore allowing researchers to infer the effects produced by presence or absence of the mycovirus under examination. Wang and colleagues observed that the presence of Trichoderma harzianum partitivirus 2 (ThPV2) did not produce negative effects on the qualitative biocontrol performance of the fungal host, which instead showed a moderate but statistically significant improved biocontrol activity in experiments with cucumber seeds inoculated with *Fusarium oxysporum* f. sp. *cucumerinum* (15). In another study, focusing on Trichoderma harzianum partitivirus 1 (ThPV1), inhibition of growth in co-cultured *Pleurotus ostreatus* and *Rhizoctonia solani* increased in ThPV1-containing strains compared with ThPV1-cured isogenic strains and this was associated to a significantly higher β-1,3-glucanase activity, whereas chitinase activity was not affected (12). Another unexpected result was obtained by You and colleagues (2019), who observed Trichoderma harzianum hypovirus 1 (ThHV1) in two forms, respectively present in two different isolates. In one of the two isolates the virus accumulates with an abundant defective form; transmission of the two versions of the infectious ThHV1 (with and without defective RNA) showed that the one with defective RNA was highly transmissible and was detrimental to the biocontrol properties of two *Trichoderma* species (16).

In this study we aimed to increase our knowledge on the diversity and distribution of mycoviruses in *Trichoderma* spp., and to better understand the possible contribution of mycoviruses to the evolution of viruses and fungi in general. We characterized the mycovirome associated with a large sub-set of *Trichoderma* spp. isolates (113 isolates) belonging to a wider collection (482 isolates) previously described by researchers from the University of Sassari: in 2009 Migheli and colleagues reported a thorough study of *Trichoderma* spp. strains isolated from 15 different soil samples, which were collected in several habitats, including undisturbed or extensively grazed grass steppes, forests, and shrub lands on the island of Sardinia, a biodiversity hotspot (19).

Here we describe a catalogue of new viruses identified in association with a sub-set of *Trichoderma* spp. isolates belonging to the aforementioned collection, that in some cases revealed previously undescribed viral clades, expanding our knowledge of virus evolution, and exploring for the first time the mycovirome associated with European populations of *Trichoderma* species.

## MATERIAL AND METHODS

### Isolates origin, growth conditions and harvesting

The fungal isolates used for this study were part of a previously described collection, hosted at the Department of Plant Protection of the University of Sassari in Sardinia and assembled in 2009 (19); our sub-set collection gathered 113 isolates of 12 different species belonging to genus *Trichoderma*, obtained from 15 different sites on the island and comprising both undisturbed and disturbed environments (forest, shrublands and undisturbed or extensively grazed grass steppes, respectively). Specific details about single isolates of our sub-set are reported in Supplemental Table S1.

Four fungal plugs obtained from monosporic cultures of each of the 113 *Trichoderma* isolates were placed on Potato Dextrose Agar (PDA) (Sigma-Aldrich, St. Louis, MO, USA) medium covered by a cellophane layer and incubated at 26 °C for 3 days in the dark. The fourth day mycelia were harvested, by gently removing each of them from the plastic layer with razorblades and placing them into 1.5 mL Eppendorf tubes along with 5 steel beads measuring 4.5 mm of diameter; lyophilization for 24 hours followed.

### Total RNA extraction

The lyophilized mycelia stored in Eppendorf tubes were disrupted by bead-beating (FastPrep-24 MP Biomedicals); the resulting powders were employed for total RNA extraction with “Spectrum Plant Total RNA” kit (Sigma-Aldrich, St. Louis, MO, USA) according to manufacturer’s protocol (20).

RNA concentration was inferred from UV absorbance at 260 nm wavelength and 260/280 ratio evaluation was used as a quality indicator: both measures were obtained using a NANODROP LITE Spectrophotometer (Thermo Fisher Scientific, Waltham, MA, USA).

### Total RNA sequencing and contigs assembly

Total RNA extracted from the Sardinian collection was sent to Macrogen inc. (Seoul, Republic of Korea) for sequencing divided in two pools, named 1 and 2 (details on pools composition are presented in supplemental Table S1). Pools were obtained by mixing 2 μg of RNA extracted from each fungal sample. rRNA-depletion and cDNA libraries were constructed using Illumina TruSeq Stranded Total RNA Gold and sequenced with a NovaSeq 6000 platform. Sequencing was then performed through “Illumina TrueSeq Stranded” approach and the resulting reads were processed by bioinformatic analysis following a well-established pipeline previously described in detail (21). Reads from RNA-Seq were first cleaned, in order to remove adapters, artifacts, and short reads through bbmap software (https://sourceforge.net/projects/bbmap/); then resulting reads were assembled *de novo* using Trinity software version 2.9.1 (22). Trinity assembly was blasted with DIAMOND software against the sorted viral portion of the NCBI nr database, and the resulting hits were manually selected and characterized through molecular approaches. The number of reads covering the viral genomes was obtained by mapping the reads from each sequenced library on virus reference sequences with Bowtie2 and read number was retrieved through SAMtools (23, 24).

Since virus identification was performed separately for each library, we compared the results of each library with the other to obtain a list of unique viruses thus reducing redundancy due to contig co-presence in both pools. Each viral contig was blasted against the whole list of viral contigs and those with nucleotide identity over 90 per cent and length over 1,000 nucleotides were grouped and considered as a single representative of the virus sequence cluster in our final list.

### ORF prediction and primer design

Starting from the above-mentioned viral contigs the respective viral open reading frames (ORFs) were predicted using the ORF finder tool from NCBI, selecting the “standard” genetic code for all contigs; primer pairs were then designed for each viral contig. Domain search on each ORF encoded by each virus was performed with “motif search” available on the GenomeNet repository (https://www.genome.jp/tools/motif/) with default parameters, along with isoelectric point and molecular weight estimation using “Compute pI/Mw” tool available on Expasy Bioinformatics resource portal (https://web.expasy.org/compute_pi/). Only ORFs with a predicted molecular mass of at least 10 kDa were graphically reported in virus genome organizations.

Primers design was performed using NCBI primer Blast tool (https://www.ncbi.nlm.nih.gov/tools/primer-blast/); results are listed in supplementary Table S2. The putative function of the predicted ORF product was suggested by Blast analysis, considering the function of the closest proteins in the NCBI database.

### cDNA synthesis and Real-time retro-transcription PCR (RT-qPCR)

Complementary DNA (cDNA) synthesis was performed on RNA extracted from each of the 113 isolates present in the Sardinian collection, using the “High-Capacity cDNA Reverse Transcription Kit” (Thermo Fisher Scientific, Waltham, MA, USA) and following the manufacturer’s protocol but halving the working volumes. Each synthesized cDNA was diluted 1:5 with sterile water for future PCR applications.

Real-Time PCR was performed using the above-mentioned primers (Table S2) on each cDNA sample according to the mapping results (i.e., in case a certain contig was present in just one RNA pool only isolates belonging to that specific pool were investigated), to associate specific viruses to specific fungal samples.

The PCR reaction was performed in 10 µL using iTaq™ Universal SYBR Green Supermix (Biorad, Hercules, USA), reaction volumes were loaded in 96-wells plates and processed by “7500 Fast Real-Time PCR system” (Thermo Fischer Scientific, Waltham, MA, USA), with thermocycling conditions of: 3 minutes at 95°C, 20 seconds at 95°C, 30 seconds at 60°C, for 35 cycles. A dissociation curve analysis was performed at the end of the RT-qPCR protocol to check for nonspecific PCR products. Isolates showing a cycle threshold (Ct) equal or lower than 31 were considered as virus-positives.

### Probe synthesis and Northern blot analysis

Infected isolates were used to clone viral genomic cDNA fragments of a length between 300 and 400 base pairs in order to obtain run off transcripts for strand specific hybridization experiments. For this purpose, cDNA was synthetized according to the previously described protocol, and then amplified in a PCR reaction using custom designed primers (Table S2). PCR products were isolated from agarose gel after electrophoresis and purified using Zymoclean Gel DNA Recovery kit (Zymo Research, Irvine, CA, USA). Purified PCR fragments were ligated in pCR® II Vector – Dual Promoter TA Cloning Kit (Invitrogen-Thermo Fisher Scientific, Waltham, MA, USA) and subsequently used for *E. coli* transformation on competent cells using Mix & Go! *E. coli* Transformation Kit (Zymo Research, Irvine, CA, USA) according to manufacturer’s protocol. Positive clones with the predicted insert (checked by digestion and subsequent vector sequencing) were used to amplify and purify plasmids; the same plasmids were linearized and used as template to synthesize DIG-labelled probes. Northern blot analysis was carried out using a glyoxal denaturation system exactly as previously described (25); DIG-labelling was obtained using “DIG RNA Labelling Mix” (Roche, Basel, CH) while for following hybridization “DIG Easy Hyb Granules hybridization solution” (Roche, Basel, CH) was employed. Signal was then detected by use of: “Anti-Digoxigenin-AP Fab fragments” (Roche, Basel, CH), “CDP-Star” solution (Roche, Basel, CH) and “Blockin Reagent” powder (Roche, Basel, CH).

### ORFan sequences detection and RNA-origin validation

Assembled contigs were submitted to a DIAMOND (v 0.9.21.122) search of the NCBI non-redundant whole database. All contigs with a homologue were discarded, whereas the remaining ones that were over 1 kb in length and encoded a protein of at least 90 amino acids (aa) (around 9 kDa) were kept, defining a preliminary set of contigs coding for ‘ORFan’ protein products. In order to select putative ORFans of viral origin alone, NGS reads were mapped on these newly obtained ORFan contigs, keeping in consideration those who mapped on both anti-sense and sense contig sequence, since a typical feature of replicating viruses is the presence of a minus and plus sense genomic template for replication (18). Further confirmation on their viral origin was later obtained by checking the absence of a DNA amplification product after PCR on total nuclei acid obtained from those isolates resulting positives for putative ORFans presence after RT-qPCR. To this extent the OneTaq® DNA Polymerase kit (New England BioLabs inc.) was adopted, exploiting above-mentioned primer pairs and following manufacturer’s protocol with a 20 µL reaction volume. Amplified bands were then separated by electrophoretic run on a TAE 1X agarose gel (1% V/V) and then UV-visualized. Positive control for DNA amplification was achieved using ITS4 and ITS5 primers (26).

### Virus names assignment

Name of viruses described in this work have been attributed using the following criteria: I) the first part of the name reflects the source of the virus (fungal species); II) the second part of the name identifies the virus taxonomical group of the first blast hit; and III) the last part of the name is a progressive number.

In case this method produced synonymous with already deposited viral sequences the progressive number was increased in order to avoid synonymous entries during sequence submission.

### Phylogenetic analysis

Genome segments encoding for RNA-dependent RNA polymerase proteins (RdRps) from all identified viruses, closest homologues, and those representatives of the phylogenetic clade present in NCBI database were aligned using the online version of Clustal Omega software, with default settings, at the EBI Web Services (27, 28). Subsequent alignment results were first screened using MEGA 11 software (https://www.megasoftware.net/) to evaluate alignment consistency of all the viral sequences under analysis, simply by checking the proper alignment of the subdomain C of the palm domain (including the GDD aminoacidic triad). Then the same alignment results have been submitted to the IQ-TREE web server to produce phylogenetic trees under maximum-likelihood model (29). The best substitution model was estimated automatically by IQ-TREE with ModelFinder (30) and ultrafast bootstrap analysis with 1,000 replicates was performed.

In addition, an estimation of the evolutionary distance between the identified viral RdRps and other viral sequences used for phylogenetic tree production has been obtained, using the ‘p-distance’ substitution model for pairwise distance computation in MEGA 11. This model expresses the proportion (p) of amino acid sites at which each pair of sequences to be compared is different. All ambiguous positions were removed for each sequence pair (‘pairwise deletion’ option). For the sake of clarity results were then transposed in aa identity, by one’s complement (1-p) using excel and thus being presented as pairwise-identity matrices.

### *In-silico* protein structure prediction and comparison

Protein structure was predicted *in-silico* using ColabFold v1.5.2 (https://colab.research.google.com/github/sokrypton/ColabFold/blob/main/AlphaFold2.ipynb)(31). Models obtained were then employed for structural comparison and Root Mean Square Deviation (RMSD) estimation using UCSF ChimeraX ‘matchmaking’ function, with default parameters. UCSF Chimera X is developed by the Resource for Biocomputing, Visualization, and Informatics at the University of California, San Francisco, with support from National Institutes of Health R01-GM129325 and the Office of Cyber Infrastructure and Computational Biology, National Institute of Allergy and Infectious Diseases (32).

### Data availability

All the raw reads generated have been deposited in the Sequence Read Archive (SRA): Bioproject PRJNA936709, Biosamples SAMN33361767 and SAMN33361768, SRR accessions SRR23531244 and SRR23531245.

## RESULTS

### Total RNA sequencing, contigs assembly and RT-qPCR

After the sequencing runs on the entire *Trichoderma* collection, we gathered a total amount of 207.195.944 reads; 105.787.432 and 101.408.512, respectively, coming from RNA pool 1 and 2. Trinity assembly generated an initial amount of 314.762 contigs; the subsequent BLAST search on a custom-prepared viral database produced a total of 26 putatively unique viral contigs (Table 1). Among those, 17 contained the typical conserved motifs of a viral RdRp, which is essential for the replication of RNA viruses and often displays three conserved aa (GDD) crucial for the catalytic activity, implying the much-likely identification of at least 17 distinct viruses. In addition, we identified 9 other segments predicted to encode for capsid proteins or protein products of unknown function and belonging to bi-or tripartite viruses. Finally, four putative viral contigs that did not show any match in nr databases were found, encoding for unknown function protein products (ORFans) without any conservation to existing catalogues through similarity searches. Among these four fragments, three (ORFan1, ORFan2 and ORFan4) actually revealed to possess RNA-only counterparts, in fact no DNA was detected corresponding to them using total nucleic acid as template for PCR amplification (see dedicated paragraph). This allowed to exclude that ORFan1, ORFan2 and ORFan4 were transcripts derived from a DNA genome, nor derived from endogenization of an RNA virus, nor that their replication occurred through a DNA intermediate.

**Table 1:**
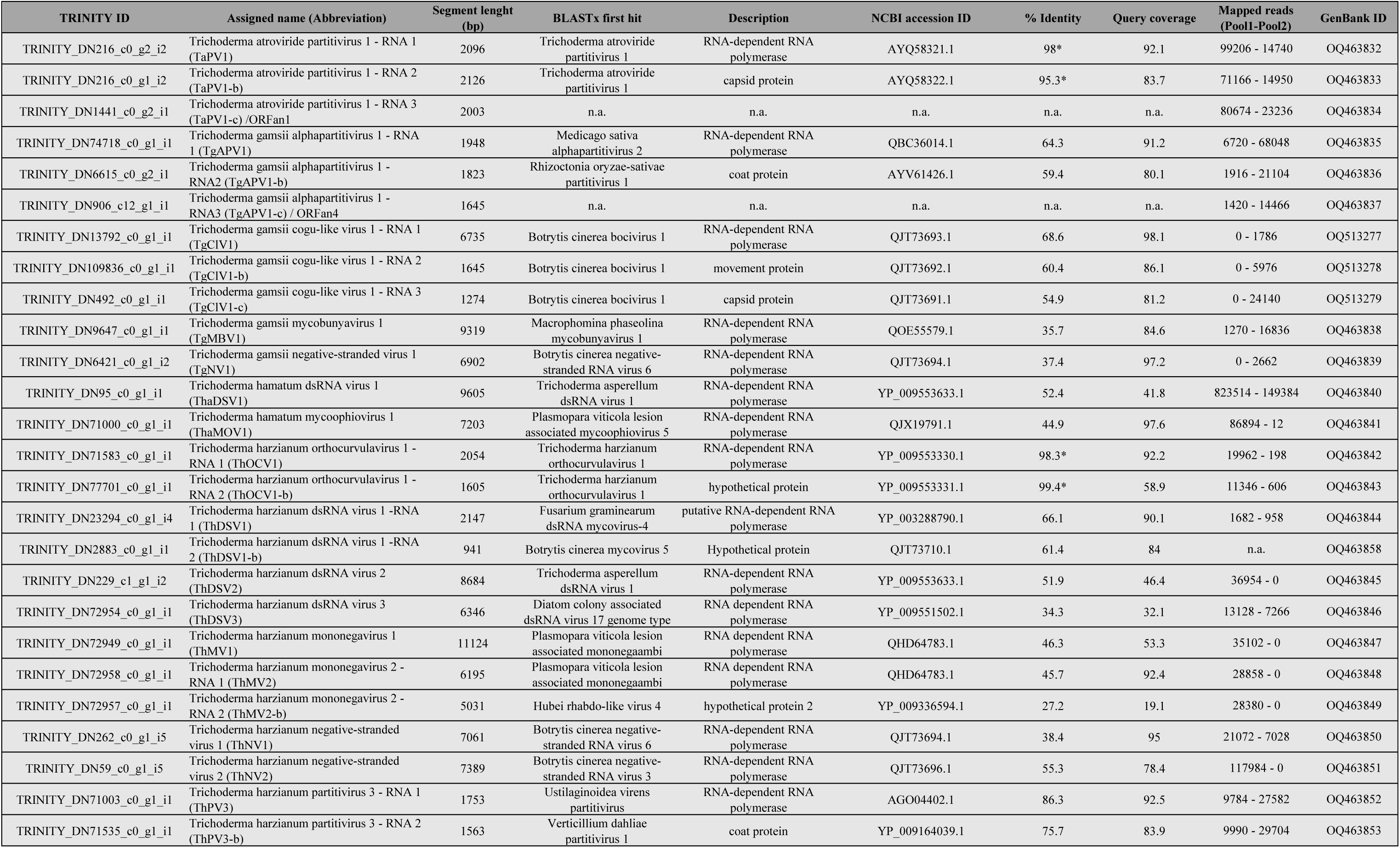
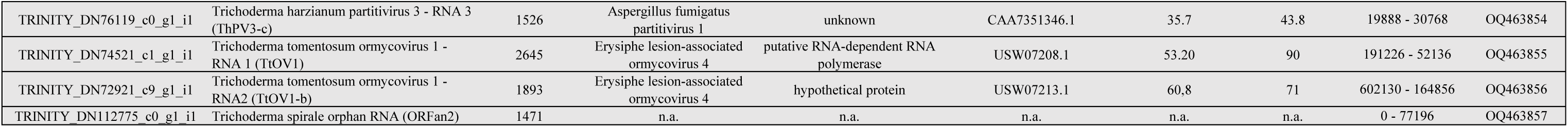
List of putative viral contigs obtained by application of our bio-informatic pipeline on rRNA-depleted total RNA extracted from Trichoderma strains. Columns show: Trinity contig ID, assigned name and abbreviation, segment length, first BLASTx hit organism, a brief description for the putatively encoded protein product of the first BLASTx hit and accession ID of the latter, identity percentage (* when >90%), query coverage and number of reads mapping for each RNA pool. N.a. stands for ‘not availabe’.

Association of each viral contig or ORFan sequence with each fungal isolate was then assessed throughout a RT-qPCR assay (Table S3, Fig. 1). The total number of different viral contigs evidenced in each single isolate is quite variable, ranging from zero to a maximum of five (e.g., isolate #99), and a quantitative estimation of the abundance of each contig can also be inferred by observing the number of mapped reads on each segment (given that concentration of total RNA from each sample in each pool was normalized). The total number of fungal isolates hosting at least one viral contig were 36 out of the 113 screened isolates; these virus-infected isolates comprised 6 fungal species belonging to: *T. harzianum* (19 isolates), *T. gamsii* (7 isolates), *T. hamatum* (6 isolates), *T. tomentosum* (2 isolates), *T. samuelsii* (33) (one isolate), *T. spirale* (one isolate) (Fig. 1).

**Fig. 1:**
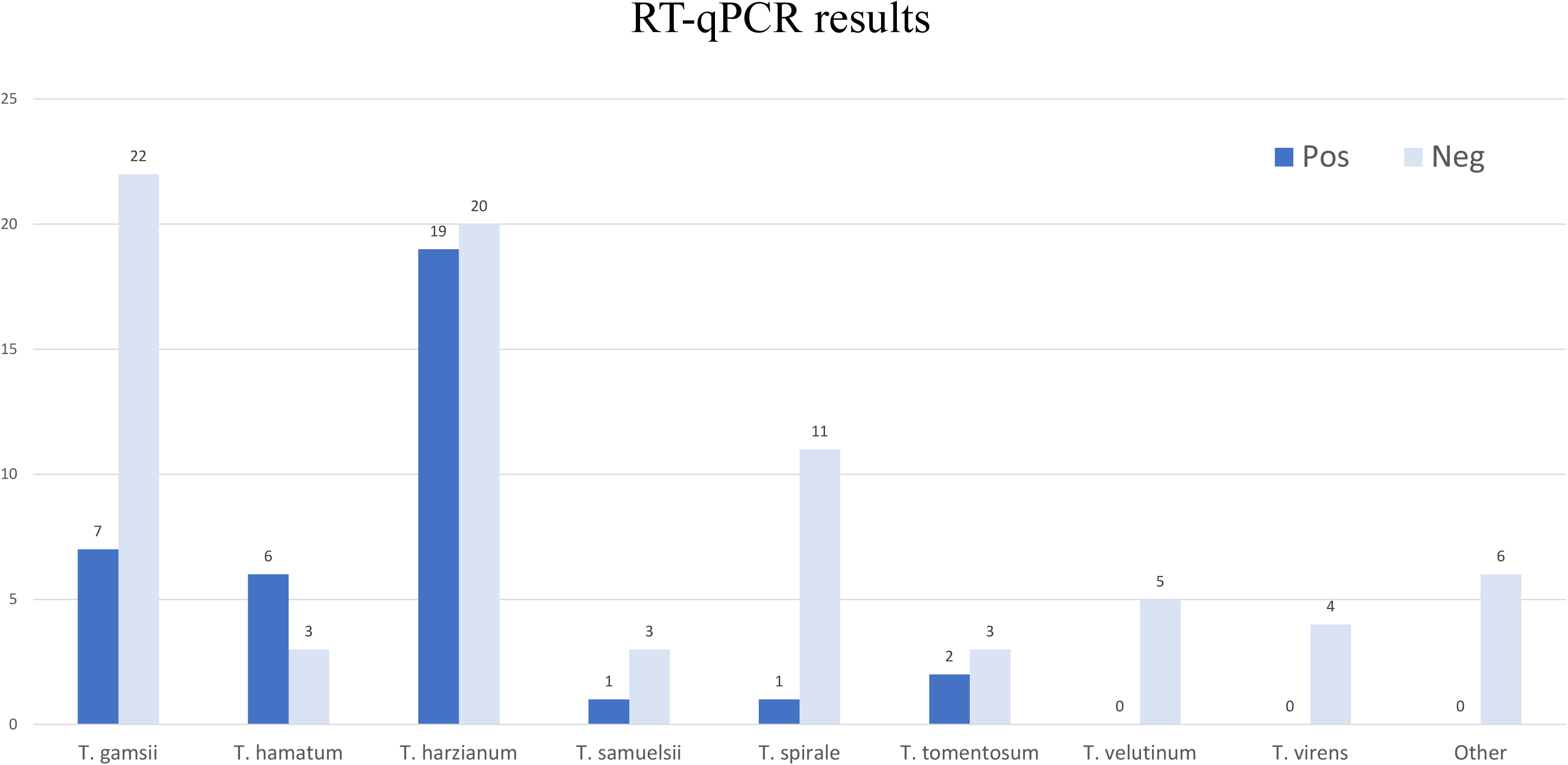
RT-qPCR results on the entire *Trichoderma* collection. Height of the bars represents the number of isolates, color represents presence (dark blue) or absence (light blue) for one or more mycoviruses characterized in this study. ‘Other’ group includes: *T. koningii* (2 isolates), *T. koningiopsis* (2 isolates), *T. asperellum* (1 isolate) and anamorph of *Hypocrea semiorbis* (1 isolate).

For bi-or multi-partite viruses, when possible, the read count/library has also been used as a guide to identify contigs belonging to the same virus; if this approach resulted in ambiguous associations, a RT-qPCR assay to find strict associations within specific isolates was used (Table S4).

The seventeen putative viral sequences encoding for an RdRp had identity ranging from 37.4% to 98.3 % to previously reported viruses. Interestingly, among those fifteen putative viral sequences were novel viruses, rather than new isolates of already known viruses; almost half of these putative viruses were predicted to have negative stranded RNA genomes and to be related to viruses present within the orders *Bunyavirales* (four novel virus sequences), *Mononegavirales* (three novel virus sequences) or *Serpentovirales* (one novel virus sequence). Additionally, another 47% of the identified viral sequences were predicted to possess a double-stranded RNA genome and have been assigned to the *Durnavirales* order (three new and two already known sequences) or to the *Ghabrivirales* order (three novel sequences). Lastly, one sequence appeared to belong to a recently proposed group of allegedly positive single-stranded RNA viruses named ‘ormycoviruses’ (34).

### Viruses characterized in the *Bunyavirales* order

Bunyavirids are mostly enveloped viruses with a genome generally consisting of three ssRNA segments (called L, M, and S) (35). The majority of the families included in this order have invertebrates, vertebrates, or plant hosts but recently specific clades infecting fungi have been characterized (36, 37). Interestingly, in our study we found only one virus (out of four) related to bunyaviruses which seems to possess a tripartite genome (i.e., Trichoderma gamsii Cogu-like Virus 1-TgClV1), while the remaining three showed only the presence of one genomic segment (Trichoderma gamsii mycobunyavirus 1-TgMBV1; Trichoderma gamsii negative-stranded virus 1-TgNV1; Trichoderma harzianum negative-stranded virus 1-ThNV1); this could be due to a lower copy number of the putative NP and other non-structural (Ns) associated proteins and/or to the fact that they are not conserved enough to be detected by homology and have escaped our ORFan detection pipeline. The L segment of these putative bunyaviruses includes the complete coding region in a single large ORF coding for the RdRp (Fig. 2). The RdRp nucleotide sequences range from ≈6.6 kb (TgClV1) to ≈9.3 kb (TgMBV1) and are predicted to encode a protein product ranging from 2200 to 3000 aa. In addition, based on ‘Motif search’ analysis, all these ORF products host a ‘bunyavirus_RdRp’ domain (pfam04196) spanning in average 400 aminoacidic residues (Fig.2, Table S5). In TgMBV1 and ThNV1 an L-protein N-terminal endonuclease domain (pfam15518) was also present on the L segment (Fig.2).

**Fig. 2:**
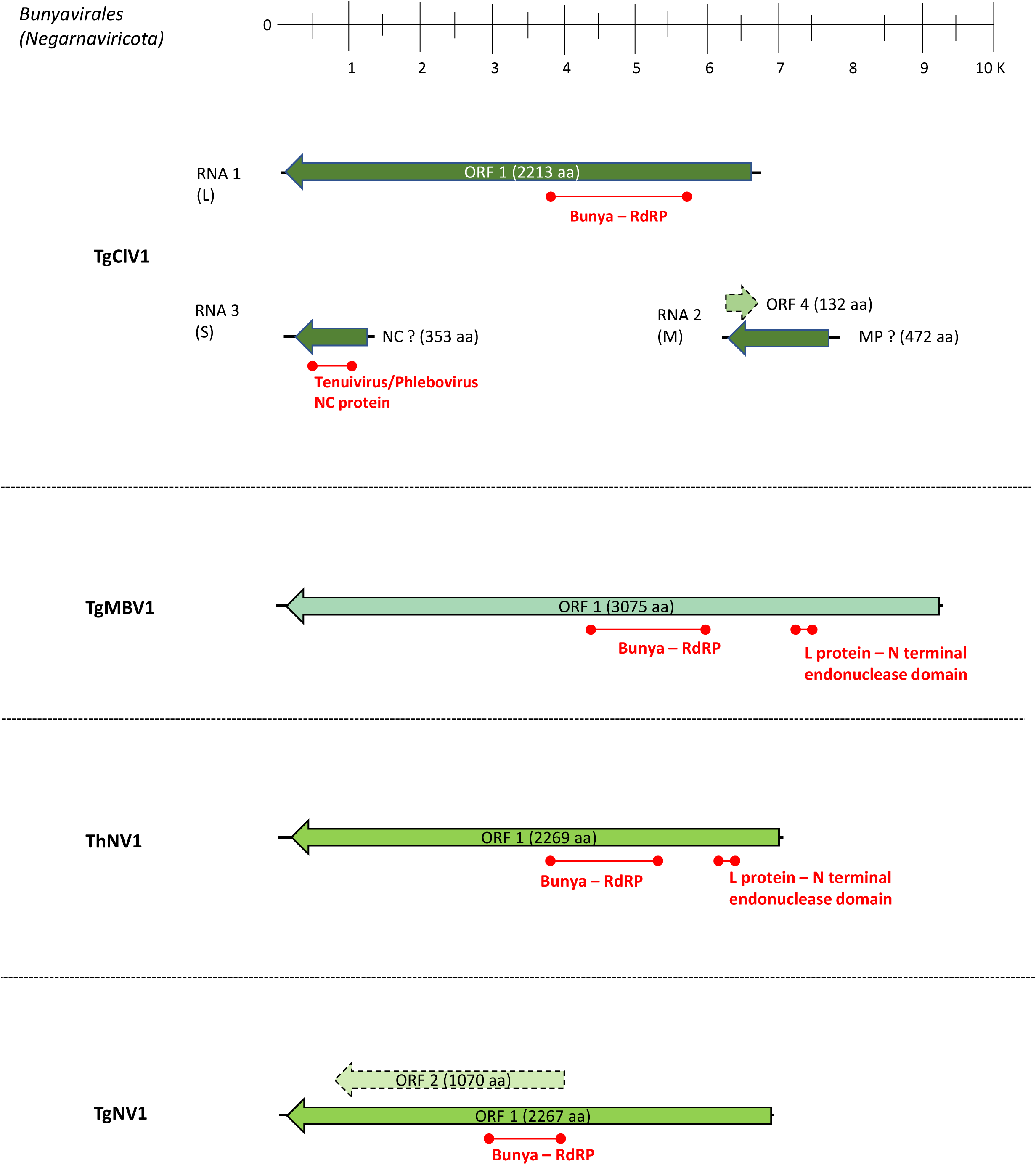
Genome organization of putative viruses belonging to the *Bunyavirales* order, top ruler indicates size in kbp. With solid lines are represented ORFs which returned at least one BlastP hit, while dotted lines represent ORFans. Presence of known protein domains predicted with Motif search analysis are highlighted in red.

For TgClV1, two other putative genomic segments were identified (RNA 2 and RNA3), respectively corresponding to putative M and S segments. RNA 3 segment is around 1300 nucleotides of length and hosts an ORF encoding for a 353 aa product having a first blast hit with Botrytis cinerea bocivirus 1-Capsid protein; Motif search analysis evidenced the additional presence of a Tennuivirus/Phlebovirus CP domain (pfam05733) (Fig. 2). RNA 2 is a segment of 1645 nt which putatively encodes for a 472 aa protein product, the latter having a first blast hit with Botrytis cinerea bocivirus 1-Movement Protein, along with an additional ORFan sequence carried on the positive strand of the segment, which should encode for an ORFan protein of 132 aa (ORF 4; Fig. 2). Phylogenetic analysis shows that all the viruses assigned to Bunyavirales based on close Blast hits are indeed in the order *Bunyavirales*, one clearly belonging to the *Phenuiviridae* family while the remaining three clustering with different unclassified *Bunyavirales* members (Fig. 3).

**Fig. 3:**
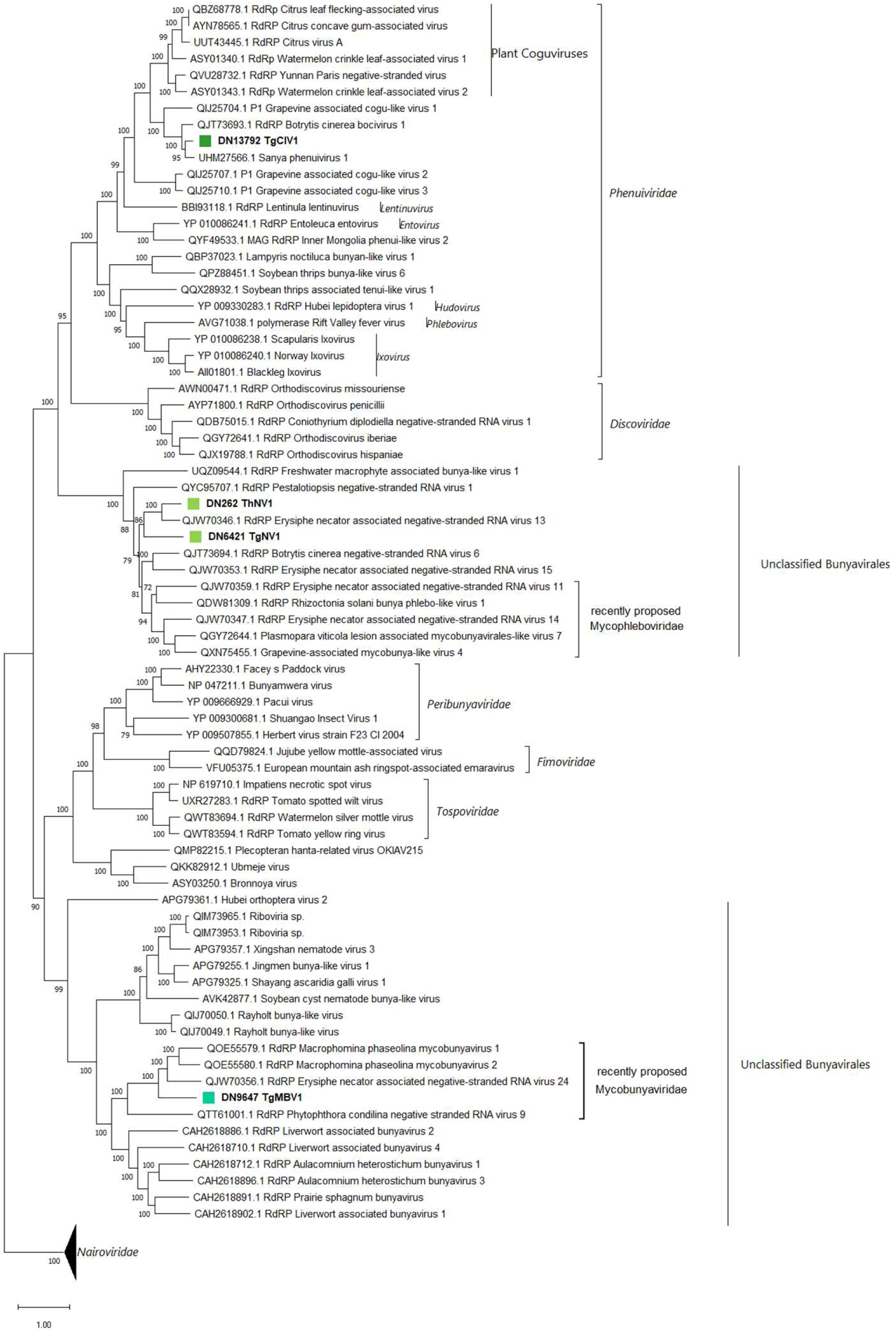
*Bunyavirales* phylogenetic tree computed by IQ-TREE stochastic algorithm to infer phylogenetic trees by maximum likelihood. Model of substitution: VT+F+I+G4. Consensus tree is constructed from 1,000 bootstrap trees. Log-likelihood of consensus tree:-396949.8245. At nodes, the percentage bootstrap values. Distinct colors indicate specific viruses in different subgroups.

TgClV1 clusters within the *Phenuiviridae* family, in a clade closely related to plant *Coguviruses*. For this reason, the name ‘Trichoderma Cogu-like virus 1’ was assigned. With respect to the remaining three viral sequences, two of them (TgNV1 and ThNV1) seem closely related to the recently proposed ‘Mycophleboviridae’ clade, which includes officially recognized mycoviruses (e.g., Rhizoctonia solani bunya phlebo-like virus 1) that, so far, have no specific Nc or NSs associated (20). Finally, according to our phylogenetic analysis, TgMBV1 seems to clearly group within the recently proposed clade of ‘Mycobunyaviridae’ (20) that includes viruses infecting fungi and oomycetes such as: Macrophomina phaseolina negative-stranded RNA virus 1 and 2, and Phytophthora condilina negative stranded RNA virus 9 (38, 39); phylogenetic analysis locate this recently proposed group of mycoviruses in a well-defined clade which seems related to viruses infecting liverwort and mosses (*Bryophyta*), and, to a lower extent, distantly related to nematodes-infecting viruses (Fig. 3). Stronger evidences, which corroborate our taxonomical hypothesis, can also be observed by examining the pairwise-identity matrices produced with MEGA 11, which express the pairwise aminoacidic identity between sequences used to build up the above-mentioned phylogenetic tree (Table S6).

### Viruses characterized in the *Mononegavirales* and *Serpentovirales* orders

The order *Mononegavirales* includes negative-stranded RNA viruses mostly monopartite, with multiple ORFs typically in the same orientation. In our analysis we have found three viruses putatively belonging to this order, namely Trichoderma harzianum negative-stranded virus 2 (ThNV2), Trichoderma harzianum mononegavirus 1 (ThMV1) and Trichoderma harzianum mononegavirus 2 (ThMV2). Initially, just two of the above-mentioned viruses were actually identified as monosegmented, while for ThMV2 a second RNA segment was detected.

The genome size of these three *Mononegavirales* members ranges from 7 to 11 kbp, and specifics can be seen in Table S5. All of them host an ORF encoding an RdRp of ≈ 1950 aa length with a characteristic catalytic domain of *Mononegavirales* RNA dependent RNA polymerase L protein (pfam00946) along with the mRNA-capping region V domain (pfam14318), known to host a specific motif, GxxTx(n)HR, which is essential for mRNA cap formation (Fig. 4).

**Fig. 4:**
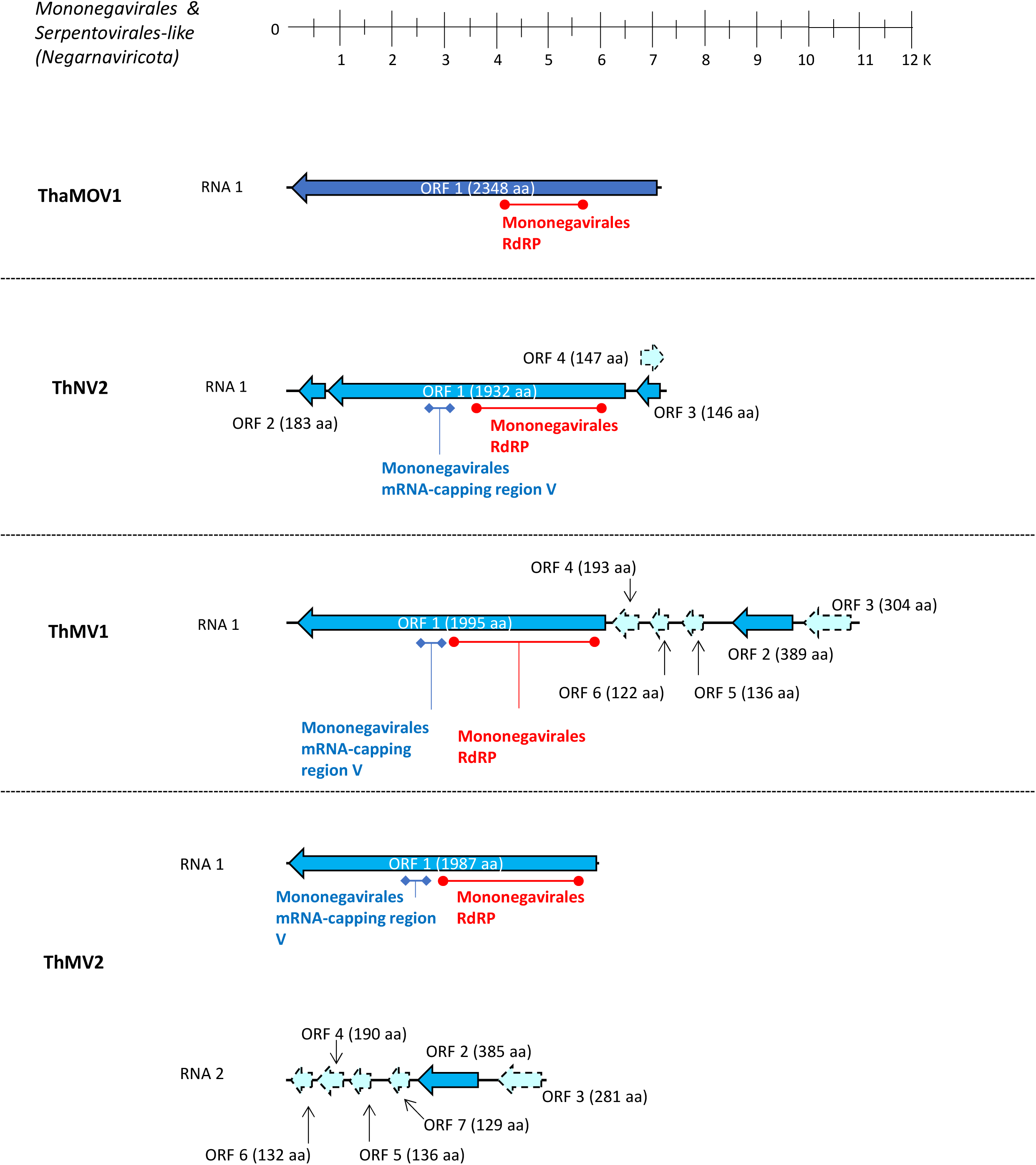
Genome organization of putative viruses belonging to the *Mononegavirales* and *Serpentovirales* order, top ruler indicates size in kbp. With solid lines are represented ORFs which returned at least one BlastP hit, while dotted lines represent ORFans. Presence of known protein domains predicted with Motif search analysis are highlighted in red and cyan.

These three putative viral RdRps had a first blast hit with negative-stranded mycoviruses, more specifically ThNV2 had as first hit Botrytis cinerea negative stranded RNA virus 3, while the other two (ThMV1 and ThMV2) both had as first hit Plasmopara viticola lesion-associated mononegaambivirus 8(21). Other additional ORFs were detected for all three *Mononegavirales* members; in facts ThNV2 hosts two ORFs flanking the RdRp-coding ORF (Fig. 4), these two small ORFs (ORF 2 and ORF 3; 552 and 441 nucleotides respectively) had a first blast hit with hypothetical proteins of unknown function belonging to Fusarium graminearum negative-stranded RNA virus 1 and Plasmopara viticola lesion-associated mononegaambivirus 2 respectively. Also, on the sense-strand of ThNV2 an ORFan coding-region was detected (ORF 4; Fig.4). With respect to ThMV1 and ThMV2-RNA 2, both possess an ORF coding for a ≈380 aminoacidic protein product as a second to last position in the reverse complement genome, which has a first blast hit with Magnaporthe oryzae mymonavirus 1 – CP (ThMV1 and ThMV2-RNA2 ORF2; Fig. 4).

Interestingly, a certain degree of synteny could be highlighted between ORFans coding regions of ThMV1 and ThMV2-RNA2 (dotted lines in Fig. 4); in facts, they all have similar size, orientation, and, to a lower extent, sequence similarity (Fig. S1, Table S7). After PCR amplification using a primer pair spanning ThMV2 RNA segments junction, no amplification product coherent with a monopartite genome organization was obtained, thus confirming the possibility of a bipartite genome organization for ThMV2. On the other hand, when performing a PCR amplification using primers specific for the same inter-genic region on ThMV1 we could obtain and visualize a coherent amplification product. Finally, after northern blot analysis we indeed confirm that ThMV1 exists as one unique genomic species of the expected size (11 kb) while ThMV2 exists as bipartite genomic species, with two segments of the expected size (6 kb for RNA1 and 5 kb for RNA2) (data not shown).

The last negative stranded RNA virus (ThMOV1) was instead hypothesized to belong to the *Serpentovirales* order, which currently include only one family (*Aspiviridae*) and one genus (*Ophiovirus*) of segmented negative-stranded RNA viruses infecting plants. We have identified the presence of one ophio-like viral RdRp in our collection; however, without detecting any presence of other associated additional segments. This single viral sequence is hosting one single ORF of ≈ 7 kbp in length coding for a putative protein of 2348 aa which again presented the characteristic catalytic domain of Mononegavirales RNA dependent RNA polymerase L protein (pfam00946) and had as first blast hit Plasmopara viticola lesion-associated mycoophiovirus 5 – RdRp; for this reason, the sequence was named Trichoderma harzianum Mycoophiovirus 1 (ThMOV1).

Phylogenetic analysis in this case suggests the accommodation of ThNV2, ThMV1 and ThMV2 within the *Mymonaviridae* family (*Mononegavirales* order); while ThNV2 directly belongs to the *Sclerotimonavirus* clade, ThMV1 and ThMV2 appear to be members of the *Plasmopamonavirus* genus (Fig. 5). On the other hand, ThMOV1 clearly belongs to a well-defined clade closely related to plant *Aspiviridae*, that includes a number of mycoviruses recently characterized, for which Chiapello and colleagues proposed the taxon name ‘Mycoaspiviridae’ (21). Pairwise-identity matrices obtained with MEGA11 (Table S8). corroborated our taxonomical hypothesis

**Fig. 5:**
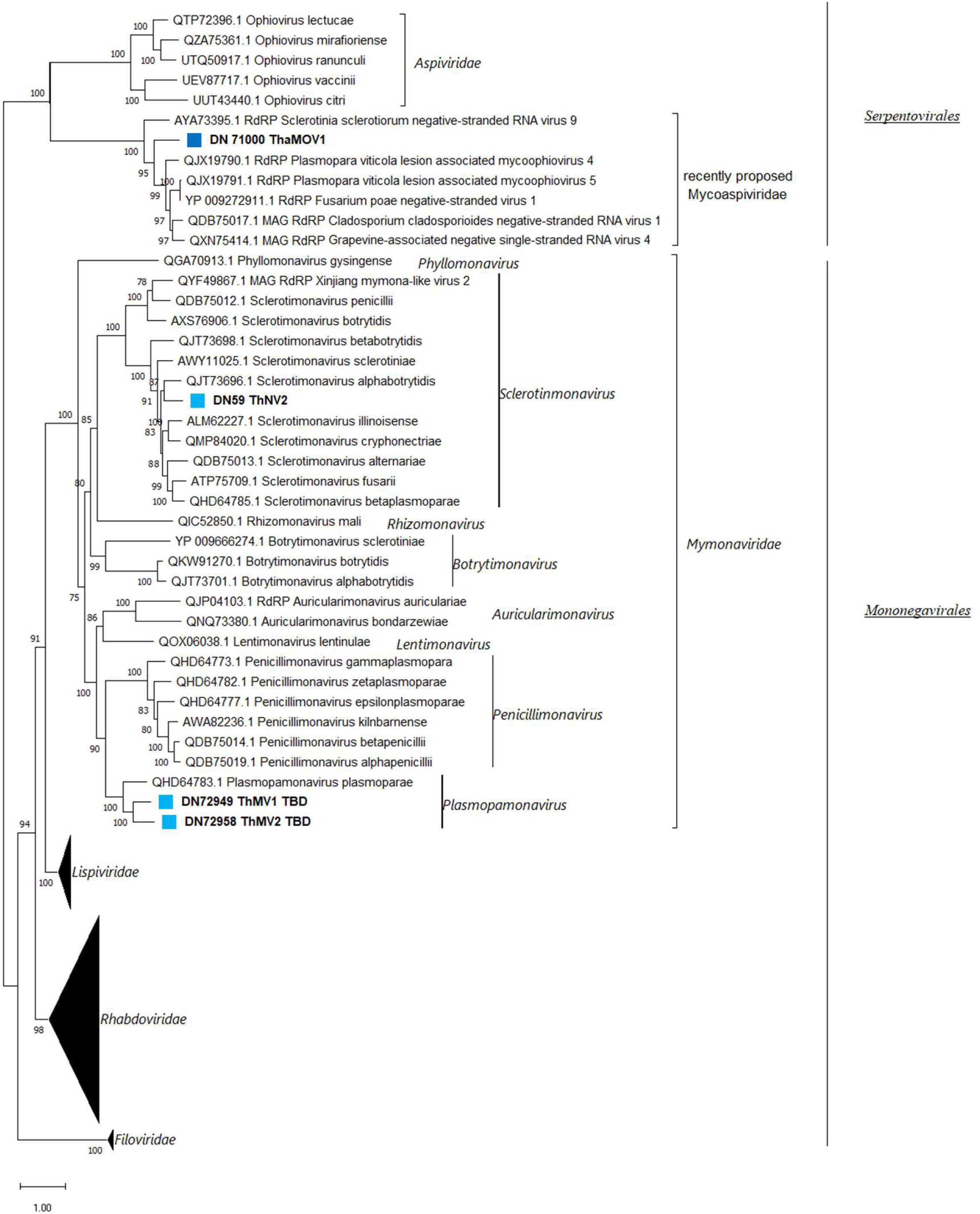
*Mononegavirales* and *Serpentovirales* phylogenetic tree computed by IQ-TREE stochastic algorithm to infer phylogenetic trees by maximum likelihood. Model of substitution: LG+F+I+G4. Consensus tree is constructed from 1,000 bootstrap trees. Log-likelihood of consensus tree:-324352.0477. At nodes, the percentage bootstrap values. Distinct colors indicate specific viruses in different subgroups.

### Bipartite *Mymonaviridae* members detected in metatranscriptomic sequences from previous studies using ThMV2 RNA2 encoded proteome as query

To shed light on the possible bipartite nature and the evolutionary origin of ThMV2, we used the putative NC sequence encoded by ThMV2-RNA2 as query for blast interrogation. This analysis was carried out on some previously described TRINITY assemblies shown to host recently characterized *Mymonaviridae* members previously included in the *Penicillimonavirus*, e *Plasmopamonavirus* genera, and reported as monopartite; some were assemblies of our property (21) while other were newly-produced ones obtained using published reads (SSR from 8303984 to 830390)(40), in both cases reads were initially originated by High-Throughput Sequencing (HTS) approaches on metagenomic samples (plant material showing grapevine downy mildew or esca disease symptoms). In this way we were able to identify 12 new contigs, undetected in the previous studies, that possessed similar size and genome organization with respect to ThMV-RNA2 (Fig. 6). Moreover, when shorts reads obtained in the two aforementioned works (21, 40) were mapped on the 11 newly identified contigs, we clearly evidenced a co-distribution of reads, shared with some previously described viral segments belonging to *Penicillimonavirus* or *Plasmopamonavirus* genus members. This evidence allows to unequivocally associate each previous RdRp-encoding segment with its newly found RNA2 (supplemental Fig. S2). All these second RNAs were therefore associated with their respective first segments, belonging to: Plasmopara viticola lesion associated mononegaambi Virus 1-2-3-4-5-6-7-8-9 (PvLamonoambi1-9), Penicillium glabrum negative-stranded RNA virus 1 (PgNSV1) and Penicillium adametzioides negative-stranded RNA virus 1 (PaNSV1).

**Fig. 6:**
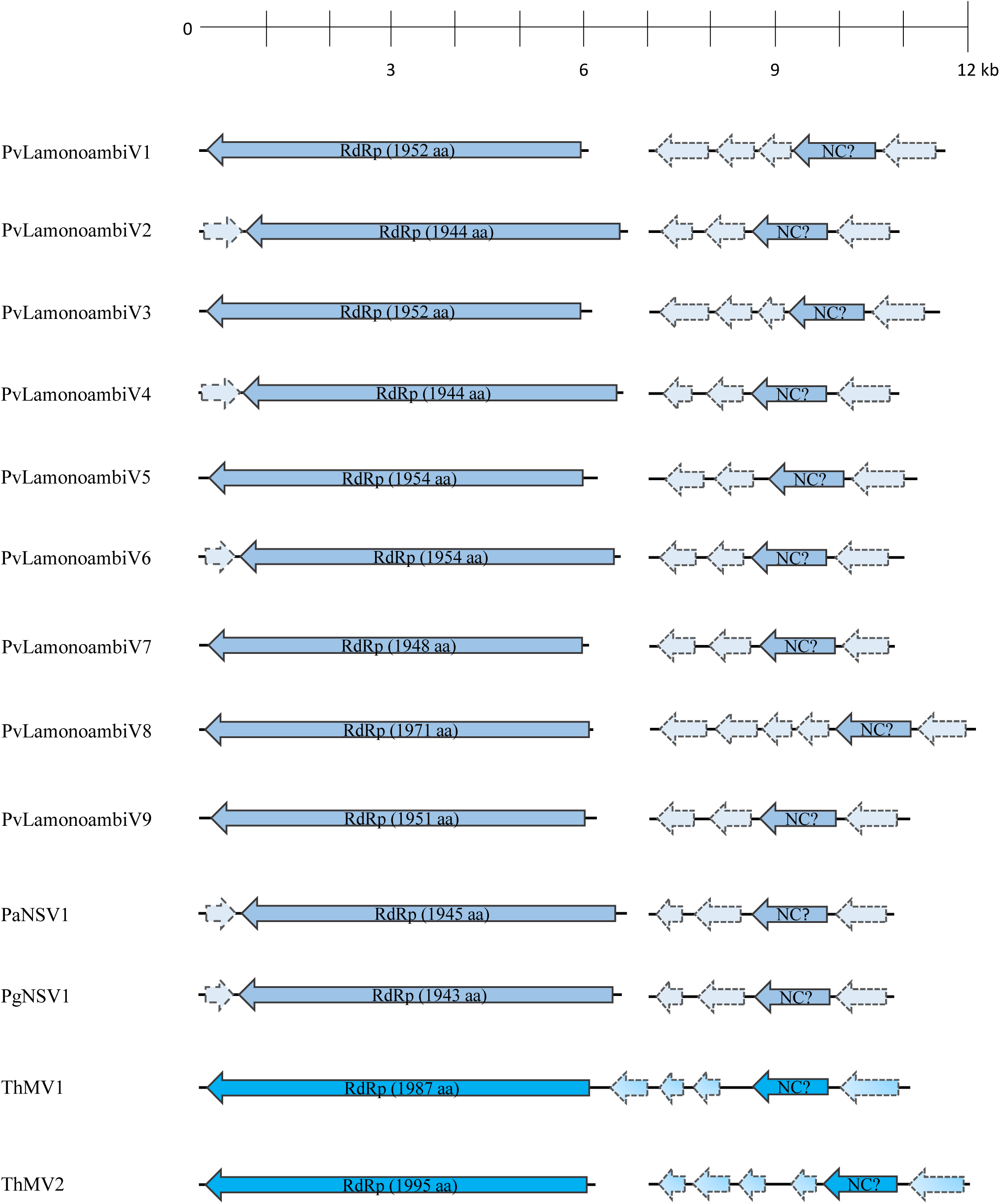
Genome organization of putative bipartite members of *Mymonaviridae* family, top ruler indicates size in kb. With solid lines are represented ORFs which returned at least one BlastP hit, while dotted lines represent ORFans. Legend: PvLamonoambiV1-9-Plasmopara viticola lesion associated mononegaambi virus1-9; PaNSV1-Penicillium adametzioides negative-stranded RNA virus 1; PgNSV1-Penicillium glabrum negative-stranded RNA virus 1; ThMV1-Trichoderma harzianum mononega virus 1; ThMV2-Trichoderma harzianum mononega virus 2.

RNA2 of the latter viruses ranged from 3900 to 5000 nucleotides in length and hosted 4 or 5 ORFs mainly coding for ORFan protein products, with the exception of the one coding for the putative NC, in second to last position of the anti-sense strand (Fig. 6). The degree of conservation evidenced by Multiple Sequence Alignment of the putative NCs led us to discard our initial hypothesis of a putative NC-coding ORF carried in sense orientation on the first RNA of some of these mymonaviruses (21), indicating that the putative NC is indeed hosted on these newly identified second segments (supplemental Fig. S3).

To further reinforce this last hypothesis, e.g. that ORF2 of RNA2 is a putative NC protein, we compared the structural conservation of these proteins (predicted in silico), with that of some experimentally confirmed NC proteins in the Mononegavirales (Sonchus Yellow Net Virus (SYNV) and Sclerotinia sclerotimonavirus (SSV)). The AlphaFold2-predicted NC protein structures of ThMV1, ThMV2, PvLamonoambiV1, PvLamonoambiV2, Sonchus Yellow Net Virus (SYNV) and Sclerotinia sclerotimonavirus (SSV) were over-imposed by matchmaking using UCSF ChimeraX, in order to highlight structurally conserved region between the latter NCs and SYNV-NC. Results of the analysis (Fig. 7) allowed the clear recognition of a structurally conserved region of two α-helices shared among all the tested NC models, spanning from residue 203 to residue 240 of SYNV (used as reference structure for matchmaking). Interestingly, this same region (203::240) indeed falls within the ‘Rhabdo_ncap_2’ Rhabdovirus nucleoprotein motif (pfam03216) exhibited by SYNV-NC in residue position 142::535 after motif search analysis.

**Fig. 7:**
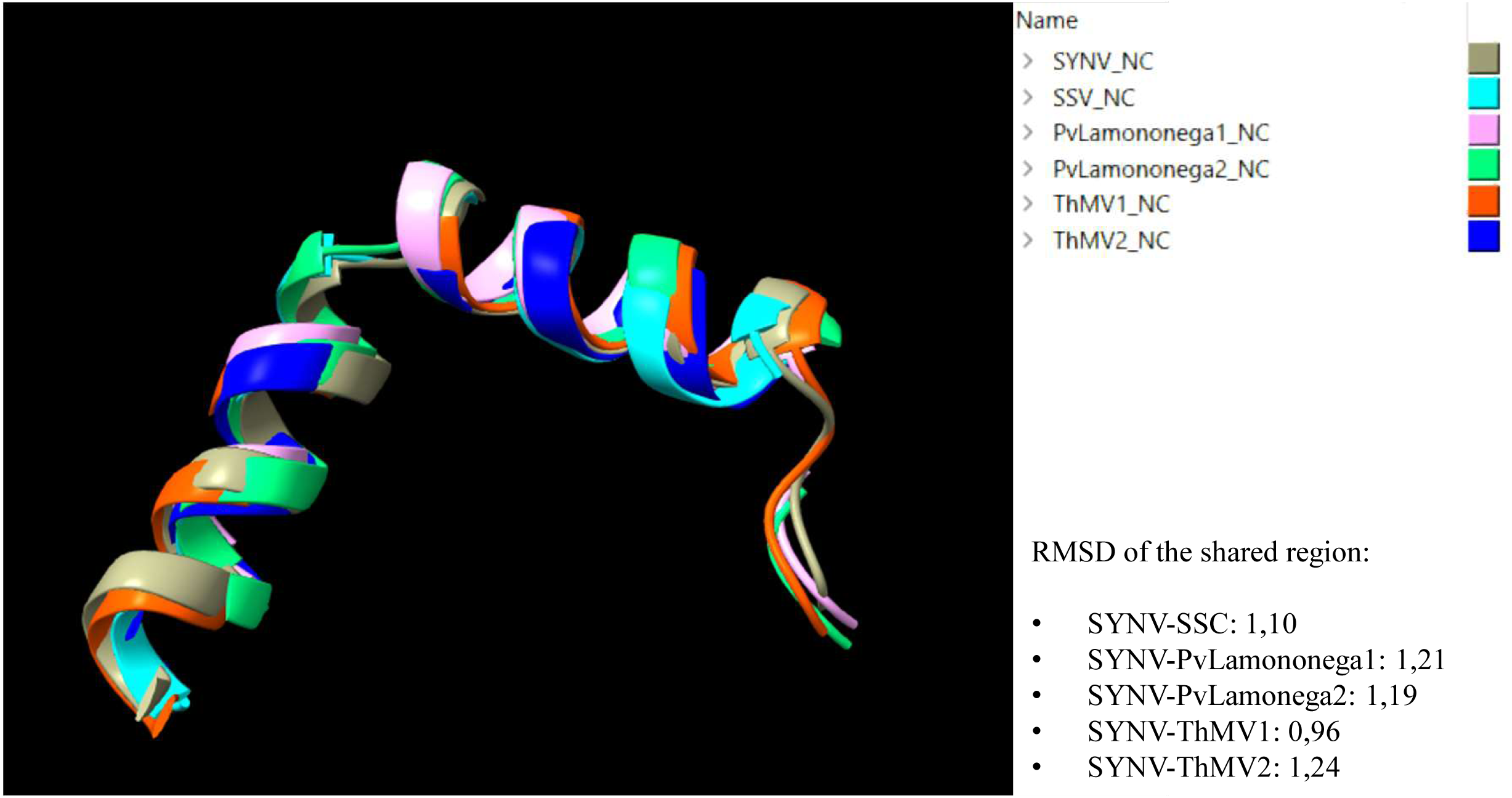
Structural comparison between NC models of different mymonaviruses (SSV, PvLamonegaambi1, PvLamonegaambi2, ThMV1 and ThMV2) and the NC model of SYNV. Different colours indicate different NCs (legend on top right of the image). Root Mean Square Deviation (RMSD) indicates the average distance in ångström between backbone atoms of different protein structures. SYNV – Sonchus Yellow Net Virus; SSV – Sclerotinia sclerotimonavirus; PvLamononega1 – Plasmopara viticola lesion associated mononegaambivirus 1; PvLamononega2 – Plasmopara viticola lesion associated mononegaambivirus 2; ThMV1 – Trichoderma harzianum mononegavirus 1; ThMV2 – Trichoderma harzianum mononegavirus 2.

### Viruses characterized in the *Durnavirales* order

Five viral sequences belonging to *Durnavirales* order were identified within our collection, among which two were already officially recognized mycoviruses (TaPV1 and ThOCV1). Three sequences were assigned to the *Partitiviridae* family, while two to the *Curvulaviridae* family. For viruses belonging to *Durnavirales* and *Ghabrivirales* orders, a finer description can be found in the Supplementary Text, along with genome representations (supplemental Fig. S4 and S6), phylogenetic trees (supplemental Fig.S5 and S7) and identity matrices (supplemental Table S9 and S10).

Interestingly, a third segment was associated to ThPV3, TaPV1 and TgAPV1; ThPV3-RNA3 hosts an ORF of 1176 bp which should encode a protein product having 35.6% identity with a protein of unknown function from Aspergillus fumigatus partitivirus 1. The third segment of TaPV1 instead, was initially referred as ORFan1 due to difficulties in association of the latter to any RdRp viral segments; nevertheless RT-qPCR analysis always detected the co-presence of TaPV1 and ORFan1 contigs (Table S3), suggesting that these three segments belonged to the same virus. The same conclusion can be applied for ORFan4 with TgAPV1, which resulted to be the third segment of the latter virus (Table S3). For this reason, we changed the name of ORFan1 to ‘TaPV1-RNA3’ and that of ORFan4 to ‘TgAPV1-RNA3’. With respect to TaPV1-RNA3, the segment carries two ORFan sequences, having opposite orientation, spanning 1605 and 291 bp (ORF 3 and 6; Fig. S4). Curiously, it is the first time that a third RNA segment is associated to TaPV1 (a virus previously characterized by Chun and colleagues (10)). Regarding TgAPV1-RNA3, it is predicted to host an ORFan sequence putatively encoding a 455 amino acidic protein product that did not return any results when subjected to blast search within nr database (Table S5; ORF3 Fig S4). The association of these third partitivirus segments to their specific partitivirus genomes is also supported by conservation of the terminal sequences (see below, ORFan results paragraph).

### Viruses characterized in the *Ghabrivirales* order

Three viruses were detected partially related to officially recognized totiviruses, but without finding any possible clear *bona fide* totivirus.

Phylogenetic analysis results clearly indicated that these novel viruses are not *bona fide* Totiviruses (member of the genus *Totivirus*); nevertheless, while ThaDSV1 and ThDSV2 seems to belong to the recently proposed family of Fusagraviridae (41), ThDSV3 clusters in a much more distant clade that includes unclassified members of the *Totiviridae* family (Fig. S7). Intriguingly, if we observe this clade of ‘Unclassified Totiviruses’, we can clearly distinguish two sub-clades: one mainly including viruses infecting sea-mosses and algae, and the other including viruses infecting invertebrates or oomycetes hosts along with ThDSV3 (Fig. S7). Pairwise-identity matrices produced with MEGA11 showed for ThDSV2 and ThaDSV1 an identity level with TaRV1 (member of this new family, Fusagraviridae) of 52.6% and 53.1%, respectively; on the other hand, ThDSV3 totalized 42.6% of identity with Prasiola crispa toti-like virus (unclassified *Totivirida*e) (Table S10).

### Ormyco-like sequences

The term “Ormycoviruses” has been recently coined to describe a novel taxonomic group including three conserved clades of protein-coding RNA segments of viral origin, typically associated with a second RNA segment with unknown function (34). Using *in-silico* structural prediction approaches, we were able to demonstrate clear structural conservation of these RNA segments with previously characterized viral RdRps, but with very limited protein sequence conservation, which initially prevented their detection through similarity searches. Moreover, within these peculiar supposed RdRps no ‘GDD’ catalytic triad was present, while the most common putative catalytic triads were ‘NDD’, ‘GDQ’ and to a less extent also ‘SDD’, ‘HDD’ and ‘ADD’ (34).

One alleged new member of this recent ‘ormycovirus’ group, which was named Trichoderma tomentosum Ormycovirus 1 (TtOV1) was identified along with a second RNA detected in strict association with the latter (TtOV1-b)(Table S4). TtOV1 segments 1 and 2 were around 2.6 kbp and 1.8 kbp of length respectively; the first is likely to encode a putative 779 a.a. long RdRp, showing 53.4% of sequence identity with Erysiphe lesion-associated ormycovirus 4 putative RdRp; while the second should encode for a hypothetical protein of 533 a.a., sharing 60.8% of identity with Erysiphe lesion-associated ormycovirus 4 (Fig. 8-A, Table S5). A reliable phylogenetic analysis that includes representatives of the five classes of RNA viruses could not be performed due to the very limited similarity of the Ormycoviruses group members to those already included in the RNA viruses’ monophyletic tree. Nevertheless, some conservation among ormycoviruses was detected through BlastX analysis by querying our ormyco-like contig (data not shown) against the whole non-redundant NCBI protein database; starting from the small sub-set (12 sequences) returned by Blast software we constructed a phylogenetic tree and a pairwise-identity matrix, just as in previous cases (Fig. 8-B, Table S11).

**Fig. 8:**
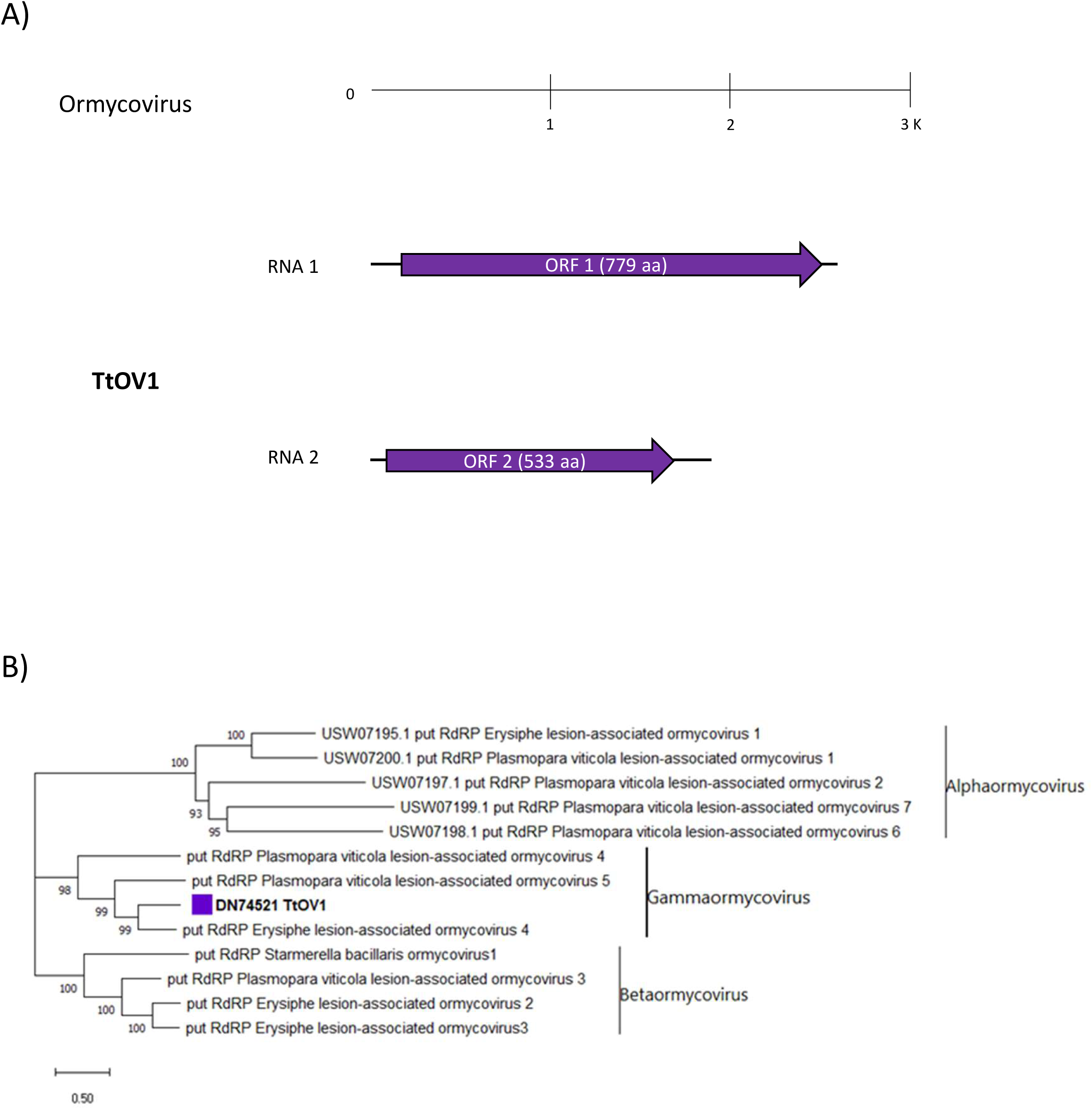
A) Genome organization of putative Ormycovirus TtOMV1, top ruler indicates size in kbp. With solid lines are represented ORFs which returned at least one BlastP hit, while dotted lines represent ORFans. B) Ormycoviruses phylogenetic tree computed by IQ-TREE stochastic algorithm to infer phylogenetic trees by maximum likelihood. Model of substitution: VT+F+G4. Consensus tree is constructed from 1,000 bootstrap trees. Log-likelihood of consensus tree:-24861.9845. At nodes, the percentage bootstrap values. Distinct colors indicate specific viruses in different subgroups.

According to our results TtOV1 is closely related to Erysiphe lesion-associated ormycovirus 4 (53.8% of identity, see Table S11) and clearly clusters within the proposed Gammaormycovirus sub-group (Fig. 8-B), further confirmation of these results came by multiple sequence alignment performed between this small sub-set of sequences (Fig. S8). TtOV1 shows the conservation of the D residue in motif A and the two G residues in motif B which are characteristic of Ormycoviruses in general, along with the catalytic triad ‘GDQ’ in motif C, known to be unique for Gammaormycoviruses (34).

### ORFan sequences

A small draft group of four ORFans, named ORFan 1, 2, 3 and 4 were identified within the collection. Starting from this group, only those having a putative viral origin (absence of amplification from total nucleic acid sample, see Fig. S9) were kept, thus reducing the number to three, namely ORFan1 ORFan2 and ORFan4; neither of the three could be located in any known viral taxonomical group and no evidence of an RdRp domain was detected within them. ORFan1 specific contig was found in association with TaPV1 contigs, thus leading us to postulate a possible role as third segment of TaPV1 genome. A confirmation of the fact that ORFan1 sequence belongs to TaPV1 genome is the degree of conservation present between the 5’ and 3’ ends of TaPV1-RNA1,-RNA2 and ORFan1 (Fig. S10); nevertheless, the ORFan nature of the protein product putatively encoded by the segment still remains unsolved. On the other hand, ORFan2 did not show any association with other viral contigs (Table S3); moreover, it was exclusively detected within one unique isolate (number #45) of *T. spirale*. ORFan2 consisted in a sequence around 1,5 kbp of length that hosted 3 putative ORFan coding sequences, each of them was averagely 260 nt long and did not return any results after ‘Motif search’ analysis (Fig. S11, Table S5). Finally, ORFan 4 could be a third segment associated to TgAPV1. In facts, again, it was found in strict association exclusively with TgAPV1 contigs; additionally, we could display some conservation on the 5’ end of the ORFan sequence with TgAPV1-RNA1 and RNA2 (Fig. S12).

## DISCUSSION

*Trichoderma* spp. are ubiquitous fungi that, as specialized saprotrophs, are able to colonize almost all environments (e.g. agricultural environment, forestry, mountain, grassland), contributing to soil and carbon mineralization. Moreover, some species are known for their potential value as biocontrol agents due to their highly-competitive mycoparasitic behaviour and their ability to improve plant health and protection, thus being broadly appreciated for agricultural applications (42). In recent years, mycoviruses have attracted increasing attention due to their effects on their hosts, but those infecting *Trichoderma* spp. have not been the subject of extensive studies; interestingly, a few available works suggest that successful application of *Trichoderma* could depend, on the long term, on presence or absence of specific viruses which, in turn, affect their mycoparasitic or antifungal activity (12, 16).

In this work we investigated the virome associated with a wide and diverse *Trichoderma* spp. collection to shed light on the diversity and distribution of mycoviruses, possibly paving the way for potential future applications within the biotechnological or agricultural domains; to this extent we chose to exploit a HTS approach on the ribosomal-RNA depleted total RNA fraction obtained from our fungal collection. This method confirmed its capability to characterize fungal viromes of vast collection of fungi, at low cost, allowing us to identify 17 viral genomes and a total of 25 viral segments (none of them endogenized in the host genome) associated to 36 out of the 113 initial different *Trichoderma* isolates evaluated. Among these 36 isolates the majority belonged to the *T. harzianum* species complex, which was the most represented group within the original collection, described in 2009 by Migheli and colleagues (19).

Interestingly, among the total number of isolates described in the latter study, the majority were positively identified as pan-European and/or pan-global *Hypocrea*/*Trichoderma* species from sections Trichoderma and Pachybasium, while only one isolate represented a new, undescribed species belonging to the Harzianum–Catoptron Clade. Moreover, internal transcribed spacer sequence (ITS) analysis revealed only one potentially endemic ITS 1 allele of *T. hamatum*, while all other species exhibited genotypes that were already present in Eurasia or in other continents (19). Evidence obtained by Migheli and collaborators pointed out to a significative decline for native *Hypocrea*/*Trichoderma* endemic populations of Sardinia, which were probably replaced by widespread invasion of species from Eurasia, Africa, and/or the Pacific Basin (19). The same isolates sampled in Sardinia were also tested for their biological control properties on a *Rhizoctonia solani*/cotton model pathosystem. A high proportion of the tested isolates (mainly belonging to *T. gamsii*) demonstrated remarkable antagonistic properties, leading to an almost complete control of the disease on artificially inoculated cotton seedlings (43).

With respect to the geographical distribution of viral sequences, the highest number of virus-infected *Trichoderma* spp. was found in soils coded F1, EG2, EG5, and EG6. The sampling sites correspond either to forest (F1) or extensively grazed (pasture) land (EG2, EG5 and EG6) (19). No correlation could be found between viral distribution and abiotic factors, such as soil properties (carbon availability), altitude, climatic conditions, and ecosystem disturbance.

This is, to our knowledge, one of the few wide-ranging studies regarding the *Trichoderma* mycovirome present in literature in term of number of isolates and diversity of species screened, along with those conducted by Jom-in (156 isolates screened), Yun (315 strains screened), and Liu (155 strains screened) (14, 44, 45). While the majority of studies on *Trichoderma* virome focused on one or few fungal strains (46), always belonging to Asian populations of the fungus, the possibility to explore such a diverse collection in a site in Europe allowed us to characterize both double-stranded and single-stranded viral strains (8 and 8, respectively) belonging to three of the major phyla that constitute the RNA viral kingdom of *Orthornavirae,* thus further increasing our general knowledge with respect to *Trichoderma* virome. This is particularly true for negative-sense viruses; in fact, to our knowledge no negative-stranded virus has been reported up to now within a *Trichoderma* host. Among the big diversity of viruses identified here, it is noteworthy the absence of members of the *Lenarviricota* phylum (mitovirus, ourmiavirus and narnaviruses) which is the most represented within ascomycetes fungi (20, 21, 47).

### A third ORFan segment associated with TaPV1

Among the 17 virus strains identified in our study, ThOCV1 and TaPV1 belonged to previously described species within the family *Curvulaviridae* and *Partitiviridae* respectively (10, 13).

In the first case, ThOCV1 was previously described by Liu and colleagues in 2019 as Trichoderma harzianum bipartite mycovirus 1 (ThBMV1) and later renamed Trichoderma harzianum orthocurvulavirus 1 (ThOCV1) (13). Among of 152 isolates belonging to different *Trichoderma* spp. collected from soil samples in China, Liu et al. (2019) identified and described one ds-RNA virus, having a bi-segmented genome, in association with only one *T. harzianum* isolate. The same viral segments (ThOCV1-RNA1 and RNA-2) have also been detected within our collection, but in association with 3 different *T. harzianum* isolates (Table S3 and Table S4).

With respect to TaPV1, Chun and colleagues published the complete bi-segmented genome of TaPV1 after total RNA extraction starting from one unique isolate of *T. atroviride* (NFCF394) collected on substrates showing green mold symptoms in a Korean shiitake farm (10). Curiously, in our analysis we have additionally identified a previously unreported third segment (TaPV1-RNA 3), the latter being present only in those fungal isolates hosting the two already known viral segments of TaPV1; this third segment was detected in all of the five isolates carrying both TaPV1-RNA1 and RNA2. TaPV1-RNA 3 segment putatively encodes for 2 ORFan protein products of 534 and 96 aa respectively (Fig. S4, Table S5), and was detected in all our TaPV1 positive samples from three distinct *Trichoderma* species.

Our hypothesis is that Chun and collaborators missed the existence of TaPV1-RNA3 because they did not investigate the possible presence of ORFan sequences within their unique *T. atroviride* isolate, just relying on similarity searches performed by blast algorithm (10). These results highlight the intrinsic limitations of methods exclusively built on similarity-based searches within existing virus databases, due to the fact that most viruses code for proteins that are not conserved enough to be detected through BLAST approaches or other profile-based methods such as HMMER (48). In this regard, we emphasize the current need of improvement for fast and reliable alternatives, for instance structural alignment algorithms (e.g., Phyre2) or targeted bioinformatic pipelines for ORFan coding segments detection (49).

Overall, the detection of TaPV1 was confirmed within isolates belonging to 3 different species present in our collection (*T. harzianum*. *T. tomentosum* and *T. hamatum*), while in the original study, Chun and colleagues found it just in one unique isolate of *T. atroviride,* collected in the Korean region of Gyonggi-do (10). These results led us to postulate that TaPV1 could possess a certain ability to overcome the species-specific barrier, potentially expanding its host range to several *Trichoderma* spp. Moreover, the possibility to detect this mycovirus within fungal isolates collected on both the Asian continent (Korea) and the European continent, could eventually reflect a long co-evolution history with the genus *Trichoderma*.

### New negative-strand RNA viruses

Negative strand viruses are a relatively recent discovery in fungi, and only a few of them have been so far characterized through virus purification and whole genome characterization (50).

In the *Bunyavirales* order, we characterized four novel viruses, but just one could be clearly assigned to an officially recognized family (TgClV1, *Phenuivirdae*), while TgMBV1 accommodated within the recently proposed Mycobunyaviridae family (20). With respect to the remaining ThNV1 and ThNV2, they appear to belong to a sister clade of the recently proposed Mycophleboviridae taxon. Given that, according to the ICTV, there are no primary classification and delimitation criteria for genus and species in the order *Bunyavirales*, pairwise sequence comparison (PASC) and phylogenetic analyses seem to be the main point of reference for new bunyaviruses taxonomy proposals; therefore, we hypothesize that ThNV1 and ThNV2 should likely warrant a new family status as soon as more members of this family are unveiled.

One of the most interesting bunyavirus found in our study was TgClV1, a tri-segmented mycovirus; each segment hosted a single ORF coding for putative Bunya-RdRp, a putative NC and a hypothetical protein with identity to the putative movement protein of Botrytis cinerea bocivirus 1 (BcBV1), a recently described negative-ssRNA cogu-like mycovirus. In their study (51), Ruiz-Padilla and colleagues describe BcBV1 as a mycovirus closely related to plant coguviruses, and, after phylogenetic analysis using both the RdRp and the NC protein, placed BcBV1 in the same clade as the plant coguviruses (51). Moreover, alignment of the core domain of the 30K viral movement protein of Laurel Lake virus (LLV), Grapevine-associated cogu-like virus 2 (GaCLV2) and Grapevine-associated cogu-like virus 3 (GaCLV3), which are currently assigned to *Laulavirus* genus, with the BcBV1 hypothetical protein showed high conservation in this region; this led the authors to suggest that this hypothetical protein could be an ancient movement protein (MP), most probably, not functional in BcBV1 (51). Since mycoviruses do not typically possess movement proteins, due to the fact that fungal hyphae have cytoplasmic continuity, Padilla and colleagues postulate a possible cross-kingdom event where an ancient plant virus, coinfecting a plant host with *B. cinerea*, was transferred from the plant host to the fungus; the resulting mycovirus BcBV1 is the product of the evolution of the ancient plant virus inside the fungus (51). In any case, when we submitted to multiple sequence alignment the aminoacidic sequence of hypothetical proteins of TgClV1 and BcBV1 along with LLV, GaCLV2, GaCLV3 and *bona fide Coguvirus* MPs (officially recognized *Coguvirus* species MPs) we could not detect any particular region showing a significative degree of conservation (data not shown). It would be of great interest to collect more reliable evidences and shed light on the possible functional role for these hypothetical proteins of BcBV1 and TgClV1; to this purpose we think that the possibility to either produce a GFP-MP fusion protein for heterologous expression *in-planta* and/or to perform a complementation assay on MP-defective phytovirus will allow to test the *in-vivo* activity of this alleged MP.

The fact that the majority of bunyaviruses characterized in this study (with the exception of TgClV1) did not possess any additional segment encoding for a NC is not surprising; in fact, numerous examples can be found in literature (21, 38, 52, 53). Besides the obvious reason linked to the intrinsic limitation of homology-based detection methods, the absence of NC coding segments could also be explained by an additional speculation. This absence could be the result of an evolutive pressure, which ensured all these bunya-like mycoviruses to obtain a modified version of their RdRp that does not need any NC, nor the common RNPs formation, to achieve a successful replication.

Regarding viruses assigned to the *Mononegavirales* order, our phylogenetic analysis clearly accommodates ThNV1 within the *Sclerotimonavirus* genus (*Mymonaviridae* family), while ThMV1 and ThMV2 within *Plasmopamonavirus* genus (*Mymonaviridae* family). ICTV taxonomy rules for members of *Mymonaviridae* (proposal code: 2020.004F) define the demarcation threshold for the species level at 80%, and for the genus level at 32% of aa sequence identity of the L protein. Consequently, we can clearly state that ThNV1 represents a novel species of sclerotimonavirus while ThMV1 and ThMV2 are novel species of *Plasmopamonavirus*.

Finally, for the only identified member of the *Serpentovirales* order (ThaMOV1) we are confident about its accommodation within the recently proposed Mycoaspiviridae family, described by Chiapello and colleagues (21).

### The first bipartite *Mononegavirales* infecting fungi

Within the order of *Mononegavirales*, members possessing a bipartite genome are quite rare and mainly belong to the *Dichorhavirus* and *Varicosavirus* genera (*Rhabdoviridae* family, *Betarhabdovirinae* subfamily). In these latter genera are included bi-segmented non-enveloped rhabdoviruses infecting plants, mainly transmitted by false spider mites (*Dichorhavirus)* or chytridiomycetes (*Varicosavirus)* (54, 55).

In this study we present strong evidence for the presence of bi-partite viruses also within the *Mymonaviridae* family, specifically within the *Plasmopamonavirus* and *Penicillimonavirus* genera. Curiously, among these newly identified bi-partite viruses, a first sub-group (PvLa1, PvLa3, PaNSV1 and PgNSV1) hosting 5 ORFs on their RNA2 could be observed, while a second sub-group includes viruses possessing only 4 ORFs on the same segment (PvLa2,-4,-5,-6,-7 and-9). For the majority of viruses present in this second sub-group, an additional ORF is instead carried in sense orientation on RNA1 and, despite all having a similar length, they do not show a strong degree of aminoacidic sequence conservation when compared to RNA2-ORF5 of the first sub-group. The remaining PvLa8 and ThMV2 instead constitutes a third sub-group which carries 6 ORFs on their second segment; interestingly the basal branch of this group consists of the monosegmented ThMV1, and therefore we have indirect evidence of a relatively recent transition from monosegmented to bisegmented genome organization.

Our collection of RNA2 from different mymonavirids allowed us to perform an *in silico* structural analysis that identified a conserved NC domain that could not be evidenced by similarity searches.

Besides a clear evidence of sequence conservation between ORFs encoding the putative NC among all these bipartite viruses (ORF2 on RNA2), it is interesting that sequence conservation can also be found when comparing ORF1 (carried in last position on the 3’-end of RNA 2) and ORF3 (in third to last position on 3’-end), but only within the three abovementioned RNA2 subgroups (i.e. those hosting 4, 5 or 6 ORFs on their second segment).

In general term, multi-partite genomes have been considered less valuable in term of transmission (cell to cell and from host to host) efficiency with respect to monopartite genomes, due to the fact that the probability of infection for each of their multiple virus particles is lower if compared with the monopartite counterparts and one or more segments could be lost during transmission (56, 57). On the other hand the hypothesis that multi-segmentation might be advantageous since it allows rapid tuning of gene expression is recently gaining acceptance (57, 58), this being particularly true after the introduction of model simulations for evaluating the effect of selective pressures applied on the genome formula (GF) (i.e. the set of genome-segment frequencies for all genome) of multipartite viruses (59). A recent study further supports the hypothesis that a possible advantage of having a partitioned genome would increase the ability to quickly modulate gene expression in highly variable environments, which require rapid adaptation responses; in fact researchers propose a scenario where the availability of a broad host-range, with widespread and well-adapted hosts in multiple environments, could contribute to drive the evolution of multipartite viruses (59). This last hypothesis finds further consistency when considering the first evidence, reported here, of a bipartite mymonavirus (ThMV2) infecting a *Trichoderma* host, a ubiquitous opportunistic fungus known to possess exceptional adaptation capabilities.

Overall, based on the previous considerations it would be reasonable to re-evaluate the taxonomic organization of family *Mymonaviridae*, taking into account the bipartite nature of some of its members.

### New double-stranded RNA viruses

Double-strand RNA viruses are currently grouped in two main lineages, due to the recent taxonomical revision of RNA viruses, namely the *Duplopiviricetes* class (phylum *Pisuviricota*), and the *Duplornaviricota* phylum (60). The fact that dsRNA viruses are not monophyletic is one of the main acquisitions from large comprehensive phylogenetic trees that includes most RNA viruses. We have identified a total of six novel ds-RNA viruses, three belonging to the *Duplopiviricetes* class (*Pisuviricota*) and three to the *Chrymotiviricetes* class (*Duplornaviricota*).

In the first case, after phylogenetic analysis and PWSA the three ds-RNA viruses (i.e., TgAPV1, ThPV3 and ThDSV1) resulted to clearly cluster within the *Alphapartitivirus*, *Gammapartitivirus* and *Orthocurvulavirus* genus, respectively, fulfilling the genus and species demarcation criteria currently adopted for the *Partitiviridae* family (61). Regarding ThDSV1, our phylogenetic analysis clearly assigns it to the *Orthocurvulavirus* genus (*Curvulaviridae* family); since this family and genus are quite recent, there are still no pre-existing criteria for official taxonomical assignment; in any case an 85% of RdRp aminoacidic identity is suggested as species demarcation criteria in the *Orthocurvulavirus* approved taxonomy proposal (number: 2020.002F).

With respect to the other group of ds-RNA viruses (*Chrymotiviricetes* class), we have identified three novel members of the *Ghabrivirales* order, named ThaDSV1, ThDSV2 and ThDSV3. Despite they could not be indicated as *bona fide Totivirus* members, our phylogenetic analysis allowed us to locate the latter of the three (ThDSV3) in a previously reported unrecognized clade of totiviruses (unclassified totiviruses-clade 1)(21, 38). Noticeably, within the latter clade we observed two distinct sub-groups, with a clear different host range: on one hand accommodating viruses mainly infecting sea-mosses and green algae and on the other hand including invertebrates-or oomycetes-infecting viruses; ThDSV3 seems to constitute an additional third sub-group which, once further knowledge will be accumulated, could eventually result as a distinct group of viruses having specific hosts, e.g., if new members infecting ‘true fungi’ are discovered. Moreover, this mycovirus was also the most common within our collection, and it was found in association with 8 isolates, belonging to 3 different *Trichoderma* spp.: *T. harzianum* (4 isolates), *T. gamsii* (3 isolates) and *T. samuelsii* (1 isolate). These results suggest that, besides TaPV1, also ThDSV3 could present a wide spectrum of host specificity, likely due to its capability of inter-and intraspecies transmission and could be quite frequent within Sardinian populations of the host.

ThDSV2 and ThaDSV1 are, to some extent, closely related to *Megabirnaviridae* and *Chrysoviridae* members (Fig. S7), yet constituting a well-supported distinct clade for which the recognition as novel family has been proposed a few years ago, under the name ‘Fusagraviridae*’* (41). According to the researchers, members of Fusagraviridae can be easily distinguished from other known mycovirus families on account of the size of their monopartite genomes (8112∼9518 bp), the genomic structure with putative Programmed –1 Ribosomal Frameshifting (–1 PRF) translational recoding mechanism (allowed by the presence of a shifty heptameric sequence, typically ‘GGAAAAC’), a long 5’-UTR (865–1310 bp) and a relatively short 3’-UTR (7–131 bp), and the arrangement of S7 and RdRp domains (41). Most of these features can be found also in our ThaDSV1 and ThDSV2 genomes, which have a coherent genome size, they both possess a putative-1 PRF motif, immediately before the stop codon UAG of the first ORF (‘GGAAAAC’ at nt 5436-5442, UAG at 5445 for ThaDSV1; ‘GGAAAAC’ at nt 4512-4518, UAG at 4521 for ThDSV2), both possess a short 3’-UTR (27 and 21 bp respectively) and both carried the ‘Viral RdRp domain’ (pfam02123) on the second ORF of the genome, like other Fusagraviridae members already described in the literature (41). The long 5’-UTR could be found only in ThaDSV1 (900 bp) while in ThDSV2 it was extremely short (33 bp), however, among some unclassified mycoviruses closely related to the Fusagraviridae group, Diplodia scrobiculata RNA virus 1 (DsRV1) and Trichoderma asperellum dsRNA virus 1 (TaMV1) also contain a short 5’-UTR of 29 and 85 bp respectively. Thus, ThDSV2 may potentially represent a further evidence for the establishment of a new genus within the Fusagraviridae family (9, 41). Finally, even if no ‘Phytoreo_S7’ (pfam07236) domain was found on the RdRp-coding ORFs of ThDSV2 and ThaDSV1 after motif search analysis, multiple alignment of RdRp aminoacidic sequences from different Fusagraviridae members highlighted a high degree of conservation within the region hosting the ‘Phytoreo_S7’ domain (Fig. S13); in conclusion we are confident that both ThaDSV1 and ThDSV2 could be considered as new mycovirus representatives of this tentatively proposed family Fusagraviridae.

### Conclusions

Results gained in this work are just an initial step towards the comprehension of the intricate mycovirome associated with *Trichoderma*, nevertheless they could contribute to further knowledge acquisition from both a plain biological standpoint and also from an agronomical and biotechnological application perspective.

Overall, taking advantage of the current progress in molecular biology, biogeography, bioinformatics, transcriptomics, proteomics and metabolomics, these studies could really contribute to, at least partially, elucidate the mechanisms underlying *Trichoderma*-mycoviruses-plant interactions. This will potentially lead to the identification of novel *Trichoderma*-based BCAs, and eventually better understand the ecology of both *Trichoderma* communities and their associated virome, within natural or artificial ecosystems.

## REFERENCES

1. Abbey JA, Percival D, Abbey, Lord, Asiedu SK, Prithiviraj B, Schilder A. 2019. Biofungicides as alternative to synthetic fungicide control of grey mould (*Botrytis cinerea*) – prospects and challenges. Biocontrol Sci Technol 29:207–228.

2. Kredics L, Antal Z, Dóczi I, Manczinger L, Kevei F, Nagy E. 2003. Clinical importance of the genus *Trichoderma*. Acta Microbiol Immunol Hung 50:105–117.

3. dos Santos UR, dos Santos JL. 2023. Trichoderma after crossing kingdoms: infections in human populations. J Toxicol Environ Heal Part B 26:97–126.

4. Zin NA, Badaluddin NA. 2020. Biological functions of *Trichoderma* spp. for agriculture applications. Ann Agric Sci 65:168–178.

5. Tripathi P, Singh PC, Mishra A, Chauhan PS, Dwivedi S, Bais RT, Tripathi RD. 2013. *Trichoderma*: A potential bioremediator for environmental clean up. Clean Technol Environ Policy 15:541–550.

6. Alias C, Bulgari D, Gobbi E. 2022. It Works! Organic-Waste-Assisted *Trichoderma* spp. Solid-State Fermentation on Agricultural Digestate. Microorganisms 10:1–12.

7. Swain H, Mukherjee AK. 2020. Host–Pathogen–Trichoderma Interaction, p. 149–165. In Sharma, AK, Sharma, P (eds.), Trichoderma, Host Pathogen Interactions and Applications, 1st ed. Springer Singapore, Singapore.

8. Ghabrial SA, Castón JR, Jiang D, Nibert ML, Suzuki N. 2015. 50-Plus Years of Fungal Viruses. Virology 479–480:356–368.

9. Lee SH, Yun S-H, Chun J, Kim D-H. 2017. Characterization of a novel dsRNA mycovirus of *Trichoderma atroviride* NFCF028. Arch Virol 162:1073–1077.

10. Chun J, Yang H-EE, Kim D-HH. 2018. Identification and molecular characterization of a novel partitivirus from *Trichoderma atroviride* NFCF394. Viruses 10:1–8.

11. Zhang T, Zeng X, Cai X, Liu H, Zeng Z. 2018. Molecular characterization of a novel double-stranded RNA mycovirus of *Trichoderma asperellum* strain JLM45-3. Arch Virol 163:3433– 3437.

12. Chun J, Yang H-E, Kim D-H. 2018. Identification of a Novel Partitivirus of *Trichoderma harzianum* NFCF319 and Evidence for the Related Antifungal Activity. Front Plant Sci 9:1–10.

13. Liu C, Li M, Redda ET, Mei J, Zhang J, Elena SF, Wu B, Jiang X. 2019. Complete nucleotide sequence of a novel mycovirus from *Trichoderma harzianum* in China. Arch Virol 164:1213– 1216.

14. Liu C, Li M, Redda ET, Mei J, Zhang J, Wu B, Jiang X. 2019. A novel double-stranded RNA mycovirus isolated from *Trichoderma harzianum*. Virol J 16:1–10.

15. Wang R, Liu C, Jiang X, Tan Z, Li H, Xu S, Zhang S, Shang Q, Deising HB, Behrens S-E, Wu B. 2022. The Newly Identified Trichoderma harzianum Partitivirus (ThPV2) Does Not Diminish Spore Production and Biocontrol Activity of Its Host. Viruses 14:1532.

16. You J, Zhou K, Liu X, Wu M, Yang L, Zhang J, Chen W, Li G. 2019. Defective RNA of a Novel Mycovirus with High Transmissibility Detrimental to Biocontrol Properties of Trichoderma spp. Microorganisms 7:507.

17. Chun J, So K-K, Ko Y-H, Kim D-H. 2022. Molecular characteristics of a novel hypovirus from *Trichoderma harzianum*. Arch Virol 167:233–238.

18. Gilbert KB, Holcomb EE, Allscheid RL, Carrington JC. 2019. Hiding in plain sight: New virus genomes discovered via a systematic analysis of fungal public transcriptomes. PLoS One 14:1– 51.

19. Migheli Q, Balmas V, Komon-Zelazowska M, Scherm B, Fiori S, Kopchinskiy AG, Kubicek CP, Druzhinina IS. 2009. Soils of a Mediterranean hot spot of biodiversity and endemism (Sardinia, Tyrrhenian Islands) are inhabited by pan-European, invasive species of Hypocrea/Trichoderma. Environ Microbiol 11:35–46.

20. Picarelli MA, Forgia M, Rivas EB, Nerva L, Carter D. 2019. Extreme Diversity of Mycoviruses Present in Isolates of *Rhizoctonia solani* AG2-2 LP From *Zoysia japonica* From Brazil. Front Cell Infect Microbiol 9:1–18.

21. Chiapello M, Rodríguez-Romero J, Ayllón MA, Turina M. 2020. Analysis of the virome associated to grapevine downy mildew lesions reveals new mycovirus lineages. Virus Evol 6:1– 18.

22. Haas BJ, Papanicolaou A, Yassour M, Grabherr M, Blood PD, Bowden J, Couger MB, Eccles D, Li B, Lieber M, MacManes MD, Ott M, Orvis J, Pochet N, Strozzi F, Weeks N, Westerman R, William T, Dewey CN, Henschel R, LeDuc RD, Friedman N, Regev A. 2013. De novo transcript sequence reconstruction from RNA-seq using the Trinity platform for reference generation and analysis. Nat Protoc 8:1494–1512.

23. Langmead B, Salzberg SL. 2012. Fast gapped-read alignment with Bowtie 2. Nat Methods 9:357– 359.

24. Li H, Handsaker B, Wysoker A, Fennell T, Ruan J, Homer N, Marth G, Abecasis G, Durbin R, Subgroup 1000 Genome Project Data Processing. 2009. The Sequence Alignment/Map format and SAMtools. Bioinformatics 25:2078–2079.

25. Rastgou M, Habibi MK, Izadpanah K, Masenga V, Milne RG, Wolf YI, Koonin E V., Turina M. 2009. Molecular characterization of the plant virus genus *Ourmiavirus* and evidence of inter-kingdom reassortment of viral genome segments as its possible route of origin. J Gen Virol 90:2525–2535.

26. Bruns TD, Lee SB, Taylor JW. 1990. Amplification and direct sequencing of fungal ribosomal RNA Genes for phylogenetics, p. 315–322. *In* PCR Protocols: A Guide to methods and Applications.

27. Sievers F, Wilm A, Dineen D, Gibson TJ, Karplus K, Li W, Lopez R, McWilliam H, Remmert M, Söding J, Thompson JD, Higgins DG. 2011. Fast, scalable generation of high-quality protein multiple sequence alignments using Clustal Omega. Mol Syst Biol 7:539.

28. Madeira F, Park Y mi, Lee J, Buso N, Gur T, Madhusoodanan N, Basutkar P, Tivey ARN, Potter SC, Finn RD, Lopez R. 2019. The EMBL-EBI search and sequence analysis tools APIs in 2019. Nucleic Acids Res 47:W636–W641.

29. Trifinopoulos J, Nguyen L-T, von Haeseler A, Minh BQ. 2016. W-IQ-TREE: a fast online phylogenetic tool for maximum likelihood analysis. Nucleic Acids Res 44:W232–W235.

30. Kalyaanamoorthy S, Minh BQ, Wong TKF, von Haeseler A, Jermiin LS. 2017. ModelFinder: fast model selection for accurate phylogenetic estimates. Nat Methods 14:587–589.

31. Mirdita M, Schütze K, Moriwaki Y, Heo L, Ovchinnikov S, Steinegger M. 2022. ColabFold: making protein folding accessible to all. Nat Methods 19:679–682.

32. Pettersen EF, Goddard TD, Huang CC, Meng EC, Couch GS, Croll TI, Morris JH, Ferrin TE. 2021. UCSF ChimeraX: Structure visualization for researchers, educators, and developers. Protein Sci 30:70–82.

33. Jaklitsch WM, Samuels GJ, Ismaiel A, Voglmayr H. 2013. Disentangling the *Trichoderma viridescens* complex. Persoonia Mol Phylogeny Evol Fungi 31:112–146.

34. Forgia M, Chiapello M, Daghino S, Pacifico D, Crucitti D, Oliva D, Ayllon M, Turina M, Turina M. 2022. Three new clades of putative viral RNA-dependent RNA polymerases with rare or unique catalytic triads discovered in libraries of ORFans from powdery mildews and the yeast of oenological interest *Starmerella bacillaris*. Virus Evol 8:veac038.

35. Ferron F, Weber F, de la Torre JC, Reguera J. 2017. Transcription and replication mechanisms of *Bunyaviridae* and *Arenaviridae* L proteins. Virus Res 234:118–134.

36. Velasco L, Arjona-Girona I, Cretazzo E, López-Herrera C. 2019. Viromes in *Xylariaceae* fungi infecting avocado in Spain. Virology 532:11–21.

37. Lin Y-H, Fujita M, Chiba S, Hyodo K, Andika IB, Suzuki N, Kondo H. 2019. Two novel fungal negative-strand RNA viruses related to mymonaviruses and phenuiviruses in the shiitake mushroom (*Lentinula edodes*). Virology 533:125–136.

38. Botella L, Jung T. 2021. Multiple Viral Infections Detected in *Phytophthora condilina* by Total and Small RNA Sequencing. Viruses 13:620.

39. Wang J, Ni Y, Liu X, Zhao H, Xiao Y, Xiao X, Li S, Liu H. 2021. Divergent RNA viruses in *Macrophomina phaseolina* exhibit potential as virocontrol agents. Virus Evol 7:1–22.

40. Nerva L, Turina M, Zanzotto A, Gardiman M, Gaiotti F, Gambino G, Chitarra W. 2019. Isolation, molecular characterization and virome analysis of culturable wood fungal endophytes in esca symptomatic and asymptomatic grapevine plants. Environ Microbiol 21:2886–2904.

41. Wang L, Zhang J, Zhang H, Qiu D, Guo L. 2016. Two novel relative double-stranded RNA mycoviruses infecting *Fusarium poae* strain SX63. Int J Mol Sci 17:1–13.

42. Mukherjee PK, Horwitz BA, Singh US, Mukherjee M, Schmoll M. 2013. Trichoderma: biology and applications. CABI.

43. Scherm B, Schmoll M, Balmas V, Kubicek CP, Migheli Q. 2009. Identification of potential marker genes for *Trichoderma harzianum* strains with high antagonistic potential against *Rhizoctonia solani* by a rapid subtraction hybridization approach. Curr Genet 55:81–91.

44. Jom-in S, Akarapisan A. 2009. Characterization of double-stranded RNA in *Trichoderma* spp. isolates in Chiang Mai province. J Agric Technol 5:261–270.

45. Yun S-H, Lee SH, So K-K, Kim J-M, Kim D-H. 2016. Incidence of diverse dsRNA mycoviruses in *Trichoderma* spp. causing green mold disease of shiitake *Lentinula edodes*. FEMS Microbiol Lett 363:fnw220.

46. Wu B, Li M, Liu C, Jiang X. 2020. The Insight of Mycovirus from *Trichoderma* spp. Agric Res Technol 24:1–4.

47. Mu F, Xie J, Cheng S, You MP, Barbetti MJ, Jia J, Wang Q, Cheng J, Fu Y, Chen T, Jiang D. 2018. Virome Characterization of a Collection of *S. sclerotiorum* from Australia. Front Microbiol 8.

48. Finn RD, Clements J, Arndt W, Miller BL, Wheeler TJ, Schreiber F, Bateman A, Eddy SR. 2015. HMMER web server: 2015 update. Nucleic Acids Res 43:W30–W38.

49. Kelley LA, Mezulis S, Yates CM, Wass MN, Sternberg MJE. 2015. The Phyre2 web portal for protein modeling, prediction and analysis. Nat Protoc 10:845–858.

50. Liu L, Xie J, Cheng J, Fu Y, Li G, Yi X, Jiang D. 2014. Fungal negative-stranded RNA virus that is related to bornaviruses and nyaviruses. Proc Natl Acad Sci 111:12205–12210.

51. Ruiz-Padilla A, Rodríguez-Romero J, Gómez-Cid I, Pacifico D, Ayllón MA. 2021. Novel Mycoviruses Discovered in the Mycovirome of a Necrotrophic Fungus. MBio 12.

52. Donaire L, Pagán I, Ayllón MA. 2016. Characterization of Botrytis cinerea negative-stranded RNA virus 1, a new mycovirus related to plant viruses, and a reconstruction of host pattern evolution in negative-sense ssRNA viruses. Virology 499:212–218.

53. Marzano S-YL, Domier LL. 2016. Novel mycoviruses discovered from metatranscriptomics survey of soybean phyllosphere phytobiomes. Virus Res 213:332–342.

54. Kondo H, Maeda T, Shirako Y, Tamada T. 2006. Orchid fleck virus is a rhabdovirus with an unusual bipartite genome. J Gen Virol 87:2413–2421.

55. Ramos-González PL, Chabi-Jesus C, Guerra-Peraza O, Tassi AD, Kitajima EW, Harakava R, Salaroli RB, Freitas-Astúa J. 2017. Citrus leprosis virus N: A new dichorhavirus causing Citrus leprosis disease. Phytopathology 107:963–976.

56. Lucía-Sanz A, Manrubia S. 2017. Multipartite viruses: adaptive trick or evolutionary treat? npj Syst Biol Appl 3:34.

57. Sicard A, Yvon M, Timchenko T, Gronenborn B, Michalakis Y, Gutierrez S, Blanc S. 2013. Gene copy number is differentially regulated in a multipartite virus. Nat Commun 4:1–8.

58. Sicard A, Michalakis Y, Gutiérrez S, Blanc S. 2016. The Strange Lifestyle of Multipartite Viruses. PLoS Pathog 12:1–19.

59. Zwart MP, Elena SF. 2020. Modeling multipartite virus evolution: The genome formula facilitates rapid adaptation to heterogeneous environments. Virus Evol 6:1–13.

60. Koonin E V., Dolja V V., Krupovic M, Varsani A, Wolf YI, Yutin N, Zerbini FM, Kuhn JH. 2020. Global Organization and Proposed Megataxonomy of the Virus World. Microbiol Mol Biol Rev 84:e00061–19.

61. Vainio EJ, Chiba S, Ghabrial SA, Maiss E, Roossinck M, Sabanadzovic S, Suzuki N, Xie J, Nibert M. 2018. ICTV Virus Taxonomy Profile: Partitiviridae. J Gen Virol 99:17–18.

